# In situ profiling of plasma cell clonality with image-based single-cell transcriptomics

**DOI:** 10.1101/2025.05.09.653118

**Authors:** Evan Yang, Jose Aceves-Salvador, Carlos Castrillon, Uli S. Herrmann, Elliot H. Akama-Garren, Michael C. Carroll, Jeffrey R. Moffitt

## Abstract

Image-based single-cell transcriptomics can identify diverse cell types within intact tissues. However, in adaptive immunity, V(D)J recombination generates unique immune receptors within cells of the same type, leading to important functional variation that is not yet defined by these methods. Here we introduce B-cell-receptor multiplexed error robust fluorescence in situ hybridization (BCR-MERFISH), which distinguishes plasma cell clones based on V-gene usage in combination with transcriptome profiling. We demonstrate that BCR-MERFISH accurately identifies V-gene usage in cell culture and in mice with restricted or native plasma cell diversity. We then use BCR-MERFISH to reveal the microbiota-dependent changes in plasma cell abundance, clonal diversity, and public clonotype usage in the mouse gut and the non-uniform distribution of plasma cell clones along the mouse ileum. As tissue context is an essential modulator of plasma cell dynamics, we anticipate that BCR-MERFISH may offer new insights into a wide range of immunological questions.

## Main

B and T cells have the remarkable ability to identify a range of pathogens and antigens due to the sequence diversity of their associated immune receptors^1,2^. B cells generate this massive sequence variation in their B-cell receptors (BCR) through V(D)J recombination and somatic hypermutation^3–6^, and, as B cells differentiate into plasma B cells, this diversity is carried over into the secreted antibody repertoire. B cells that carry highly similar BCR sequences are considered clones, and changes in B cell clonality within a tissue are indicative of B cell immune responses^7–9^. Sequencing-based methods such as bulk or single-cell BCR-seq have proven adept at characterizing changes in clonality and have been used to link high clonal diversity to proper immune surveillance and tissue homeostasis, or low clonal diversity to infection^10–12^, autoimmunity^12–14^, and cancer^15^.

While the B cell response can act systemically through circulating antibody-secreting plasma B cells, many B cell behaviors are inherently local and involve specific cellular interactions in different tissue contexts. For example, within germinal centers local interactions between T follicular helper cells, follicular dendritic cells, and unique germinal center B cell clones regulate the selection and clonal expansion of specific clones^16–18^. Similarly, local interactions are also important in disease. For instance, aberrant interactions between B cells and T follicular helper or dendritic cells has been implicated in the breakage of immune tolerance and the production of autoantibodies^16,19–21^. More broadly, atypical accumulation of B cells in specific tissues or tissue sites has been associated with diseases such as ulcerative colitis^22^, B cell cancers^23,24^, and systemic lupus erythematosus^13,25^.

The importance of cellular interactions and tissue context in these B cell functions has led to the introduction of technologies that can provide spatially resolved measurements of B cell clonality. For example, Confetti mice discriminate B cell clones through one of ten different genetically encoded two-fluorophore combinations and have been used to reveal the evolution of germinal center (GC) B cells towards pauciclonality within autoreactive GC of the spleen^25^ or in Peyer’s Patches (PP) following oral immunization^26^. Yet as a genetically encoded fluorescent measure of clonality, Confetti provides no direct link to sequence features of the BCR, provides limited information on the cellular identity and behavior of surrounding cells, and is restricted to genetically modified organisms. Recently spatial transcriptomic methods have been extended to characterize features of the adaptive immune system. Spatial V(D)J^27^, SPTCR-seq^28^, Stereo-XCR-seq^29^, and Slide-TCR-Seq^30^ leverage spatial capture arrays to characterize both transcriptome expression and sequence features of the BCR^27,29^ or the T cell receptor (TCR)^28,29^ and, thus, reveal the structure of tissues via gene expression, provide an estimate of local cellular content, and define local measures of B-cell or T-cell clonality. However, the resolution of these methods is multi-cellular, either due to physical limits on capture resolution (10 to 50 microns^27,28,30^) or the need to spatially aggregate sparse RNA measurements^29^; thus, these methods have only reported an inference of the local environment around immune cells or of the single-cell linkage of heavy and light chains^27–30^. For these reasons, there remains a need for spatially resolved methods that can directly characterize sequence features of immune receptors while simultaneously profiling gene expression, both at single-cell resolution.

Here, we present BCR-MERFISH (B Cell Receptor Multiplexed Error-Robust Fluorescence In Situ Hybridization), a single-cell spatial method that simultaneously measures plasma B cell clonality and the transcriptome of their surrounding tissues (Fig. 1a). BCR-MERFISH is built on MERFISH^31,32^, which leverages successive rounds of fluorescence in situ hybridization (FISH) to create optical barcodes that distinguish hundreds to thousands of unique RNA molecules with sub-cellular resolution within intact tissue samples^31–36^. In BCR-MERFISH, B cell clones are identified via optical barcodes that reveal the unique combination of heavy and light chain V gene usage as well as the heavy chain constant region. To overcome sequence homology between V regions, we introduce homology-aware probe design and barcoding schemes that leverage the highly expressed BCR mRNA in plasma cells. We demonstrate accurate V gene identification in cell culture, in mice with a restricted BCR repertoire, and by comparing to BCR-seq of wild type mice. To demonstrate the biological potential of BCR-MERFISH, we chart the abundance, location, and clonality of plasma cells in the gut of specific pathogen free (SPF) or germ-free (GF) mice. These measurements demonstrate the role that the microbiome plays in plasma cell abundance and clonal diversity, confirms that public clonotypes seen in GC B cells are represented in plasma cell populations, and discover that plasma cells are not uniformly distributed even along short lengths of the small intestine. With the ability to simultaneously define plasma cell clonality based on sequence features of the BCR and map cell types and states in the surrounding tissue, we anticipate that BCR-MERFISH may prove to be a useful tool for profiling interactions with and effects of elements of the adaptive immune system in a wide range of tissue contexts.

**Fig. 1.**
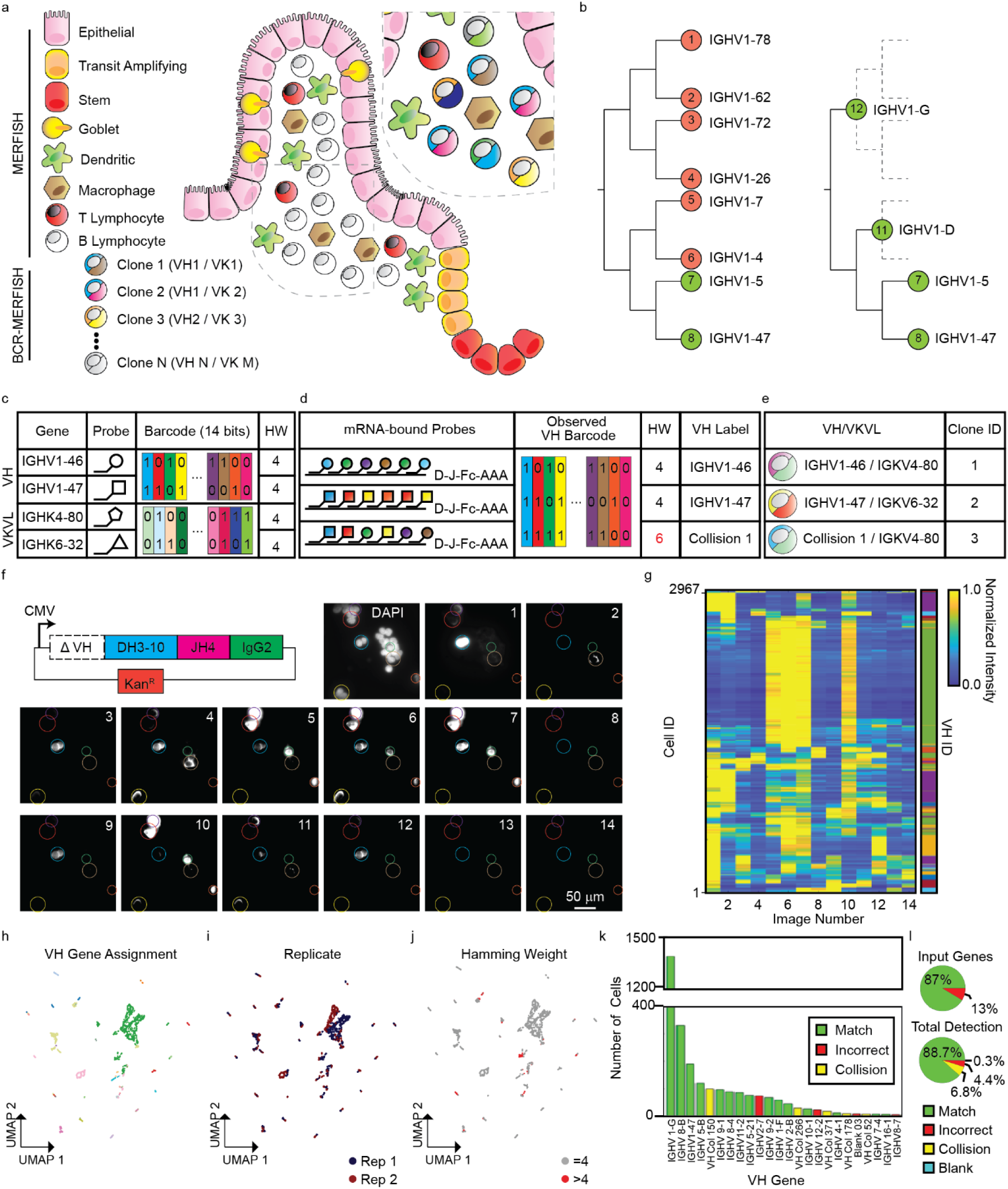
Homology-aware probe design and homology-resistant barcoding allow identification of sequence features of the B cell receptor. **a**, MERFISH defines cell types and states with gene expression while BCR-MERFISH defines B cell clonality through the co-expression of heavy and light chain V genes. **b**, Illustration of how homology-aware probe design manages high degrees of V-gene similarity by grouping homologous subsets. Green and red indicates V genes or V gene groups for which sufficient probes can or cannot be designed, respectively. **c**-**e**, Illustration of a homology-resistant encoding scheme. Different V genes are assigned barcodes with a fixed Hamming Weight (HW; c). V genes that bind probes targeted to other V genes will generate barcodes with a greater HW, indicating a collision (d). Plasma cell clones are defined by the pairing of VH and VKVL genes, even those defined as a collision (e). **f**, Expression plasmid for a heavy chain with different VH choices (top) and the fourteen images used to decode the VH gene within HEK293 transfected with a mixture of VH plasmids (bottom). A DAPI image highlights the location of all cells, and circles highlight cells expressing the plasmid with color indicating different optical barcodes. Scale bar: 50 µm. **g**, Normalized intensity (left) for each transfected cell across all images sorted by assigned VH ID (right, color). **h-j**, UMAP representation of the intensity profiles for each cell in (g) colored by the VH ID (h), experimental replicate (i), or barcode HW (j). **k**, Histogram of the number of cells assigned to each VH ID sorted by abundance. Color indicates identified VH genes included (green, Match), not included (red, Incorrect), or which were identified as a collision (yellow, Collision). **i**, Fraction of included or not included VH genes detected (top) or fraction of all cells identified as an included VH (Match), a not included VH (Incorrect), a collision barcode, or as one of the false-positive control barcodes (Blank).

## Results

### Identifying B cell clones with homology-aware probe design and encoding

The central challenge to identifying B cell clones via the sequence of the BCR with FISH-based methods is the high degree of sequence homology within the gene segments that comprise the heavy and light chains of the BCR. During B cell maturation, the primary sequence of the heavy and light chain of the BCR is created through V(D)J recombination^5,7,37^. The many tens of germline V, D, or J gene segments from which this process selects arose through genetic duplication^38^ and therefore share high degrees of sequence homology. Unfortunately, homologous genes are challenging for FISH-based detection, as hybridization permits some sequence mismatch; thus, probes designed for one RNA target may bind several off-target RNAs with sufficient homology. Standard FISH-probe design approaches identify and discard such probes^32^; however, for sets of genes with substantial degrees of homology, these design approaches may produce no viable probes for many targets.

Here we address this limitation by introducing two complementary approaches. First, we developed a homology-aware FISH-probe design algorithm (Methods). The key insight to this approach is that sets of homologous genes can be organized via their relative similarity and that homology between gene subsets can be managed by targeting groups of highly similar RNAs, rather than individual RNAs (Fig. 1b). Specifically, we developed an iterative algorithm that leverages a similarity-tree-based representation of relative homology to guide probe design. In this approach, standard probe design methods^32^ are used to identify all target RNAs with sufficient dissimilarity from all other targets to support a desired number of unique probes per RNA. Any RNA that does not support sufficient probe numbers—because of its high degree of homology to other RNA—is then grouped with the nearest RNA or set of RNAs, as determined by the similarity tree. Probe design is then repeated with the critical difference that potential homology to other RNAs is only considered for RNAs outside of a group and potential probes are not penalized for homology to RNAs within a group. This process then iterates until enough probes have been designed for all RNAs or groups of RNAs (Fig. 1b).

We used this homology-aware probe-design approach to group all 188 IGHV (VH) genes and 102 IGKV/IGLV (VKVL) into 80 VH and 68 VKVL targetable groups (Fig. S1a-d, Table S1, Methods). We did not use this approach to design probes against the D or J segments due to their shorter length. Despite their high degrees of homology, 67 of the 188 VH genes were directly targetable with the remaining genes organized into 13 groups of various sizes (Fig. S1a,b; Table S1). Similarly, 63 of the 102 VKVL genes were directly targetable with the remaining genes organized into 5 groups (Fig. S1c,d; Table S1). Supporting our analysis, our similarity trees reproduced the established family organization for both the heavy and light V genes (Fig. S1a,c) and all members of each of the V gene groups produced by our algorithm were drawn from the same families (Fig. S1a-d; Table S1). For genes that could be detected without grouping, we chose to label them by their gene name (e.g., IGHV1-47) whereas grouped genes were named via the gene family that comprised that group followed by a unique letter (e.g., IGHV1-A or IGHV1-B; Table S1).

Despite this grouping of high homology genes, we reasoned that some degree of off-target binding between groups may still occur due, perhaps, to residual homology or sequence modification by somatic hyper mutation. Thus, we also created a homology-resistant encoding scheme (Fig. 1c-e). In this approach, targetable genes or groups are assigned a unique binary barcode with a fixed Hamming weight (HW), i.e., the number of bits with a value of ‘1’ (Fig. 1c). All RNAs that are bound by only probes targeted to that RNA would generate a barcode with that fixed HW (Fig. 1d). By contrast, if an RNA had bound probes from multiple targets, due to residual homology between the two, it would generate a signal that represents the combination of both barcodes—an event we call a collision—which would be clearly detected by a larger observed HW (Fig. 1d). Such approaches have been used previously with MERFISH to identify and discard error containing barcodes^31^; however, as collisions are driven by the sequences of the V genes, in this case, collision barcodes are likely V gene specific and, thus, could still be used to distinguish B cell clones (Fig. 1e). However, as it is not possible to uniquely determine the barcodes that collided because such barcodes cannot be associated with unique V gene groups; thus, we label these unique collision events separately (i.e., Collision 1; Fig. 1d,e; Methods; Table S1).

Leveraging these approaches, we generated a set of MERFISH encoding probes that collectively define the different VH and VKVL groups. VH and VKVL genes are decoded by separate 14-bit barcodes, each associated with an orthogonal set of readout probes to allow their combined measurements (Tables S1&S2; Fig. S1e-j). In addition, as B cells also undergo class switch recombination^39^ to select a specific constant region of the heavy chain (Fc), we also generated probes against the 7 different Fc regions (IgD, IgM, IgA, IgE, IgG1, IgG2, and IgG3; Tables S2, S3, &S4). Collectively, these probes should be able to distinguish between 80×68=5440 clones defined by their unique VH/VKVL pairs. Additionally, with the 7 different Fc regions, we can distinguish 5440×7=38,080 different possible B cells (Fig. S1e-j). Nonetheless, it is important to note that this definition of clonality differs from that used by sequencing-based methods, which typically includes both the selection of V, D, and J region as well as some degree of homology (often 90%) in the sequence of the complement determining region 3 (CDR3)^40^. Finally, due to unique readout probes being associated with each bit, these measurements can be performed in combination with standard MERFISH, allowing plasma cell clonality tracking to be layered upon standard MERFISH measurements (Fig. 1a). Collectively, we term this approach BCR-MERFISH.

### BCR-MERFISH accurately identifies VH and VKVL gene groups in single cells

To test the accuracy of V gene labeling with our homology-aware and -resistant probes, we created high expression plasmids containing a variety of distinct VH genes in combination with a defined D and J, driven by the constitutively active Cytomegalovirus (CMV) promoter (Fig. 1f; Methods). We then transfected HEK293 cells with a set of plasmids containing a known subset of VH genes and imaged them with BCR-MERFISH using probes targeting only the VH genes (Methods). As expected, individual cells showed bright fluorescent signals that varied in their on-off pattern of fluorescence across the multiple rounds of imaging, with all transfected cells showing the same pattern when a single construct was used and different cells showing different patterns when a complex collection of plasmids were used (Fig. S2a,b; Fig. 1f).

To quantify the patterns of observed fluorescence, we integrated the fluorescence within individual cells and normalized the intensities across imaging rounds (Fig. 1g; Fig. S2c). In samples containing multiple different VH genes, we observed a diversity of optical patterns that were statistically distinguishable (Fig. 1h), were reproduced across replicate measurements (Fig. 1i), and showed limited numbers of patterns that failed our collision detection (Fig. 1j). Importantly, essentially all patterns that passed the collision detection could be matched to barcodes associated with individual VH groups (Fig. 1g,h; Methods), cells assigned to the same VH group showed highly consistent optical barcodes (Fig. S2d), and VH groups that contained more V genes, such as IGHV1-G, were observed more frequently than V gene groups associated with a single V gene (Fig. 1h,k; Fig. S2d). Collectively, these observations support the ability of BCR-MERFISH to distinguish VH gene groups.

To quantify the accuracy of BCR-MERFISH, we compared the identified VH gene groups to the subset of V genes included in these measurements. 87% of VH genes included in our plasmid set were detected with BCR-MERFISH in at least one cell, while 89% of transfected cells were matched to a V gene included in the set (Fig. 1k,l), supporting the accuracy of BCR-MERFISH. Moreover, of the distinguishable optical patterns we observed, only 7% were detected as collision events, suggesting both that our probe design strategy was effective at managing the high-degree of homology between V genes and that our collision-detection scheme could identify these cases. Finally, we included in our barcode set valid barcodes that were not assigned to individual V genes and for which no encoding probes were created. These ‘Blank’ barcodes serve as an internal measure of our false positive rate (Table S1). Only 0.3% of cells were associated with a Blank barcode, providing additional support for the accuracy of our V gene calling with BCR-MERFISH.

We performed similar measurements with the VKVL genes and observed a similar accuracy in our ability to define VKVL with BCR-MERFISH (Fig. S2e-h). Taken together, these experiments support the ability of BCR-MERFISH to both distinguish distinct heavy and light chain V gene groups with optical barcodes and, in the vast majority of cases, associate these unique barcodes with the identity of the V gene group.

### BCR-MERFISH accurately identifies V gene groups in mice

In order to validate the ability of BCR-MERFISH to identify V gene groups *in situ*, we first turned to a transgenic mouse that harbors a knock-in of a prearranged BCR (564Igi; C57BL/6)^25,41,42^. As this mouse is RAG-proficient and contains an unarranged BCR locus, we anticipate some diversity in V gene usage. Nonetheless, we expect a strong enrichment of plasma cells using the prearranged heavy and light chain V gene pair (564H: IGHV1-53, 564K: IGKV4-59; Fig. 2a)^25,41,42^. To determine whether BCR-MERFISH observed this enrichment, we harvested a tissue rich in plasma cells—the ileum—from 8-week-old 564Igi mice and imaged sections of this tissue with BCR-MERFISH (Fig. 2b; Methods). As expected, we observed IgA+ plasma cells in the lamina propria (LP) of these ileal slices (Fig. 2b). Importantly, BCR-MERFISH identified the expected limited diversity of VH and VKVL gene groups within these plasma cells (Fig. S2i,j): 53% of plasma cells were associated with 564H and 96% with 564K. Moreover, we generated single-molecule FISH probes (Table S2) targeting only the 564H and 564K V genes and found that they further supported the presence of IgA+/564K+ ileal plasma cells that do not use the 564H V gene (Fig. S2k). Collectively, these measurements support the *in situ* accuracy of BCR-MERFISH.

**Fig. 2.**
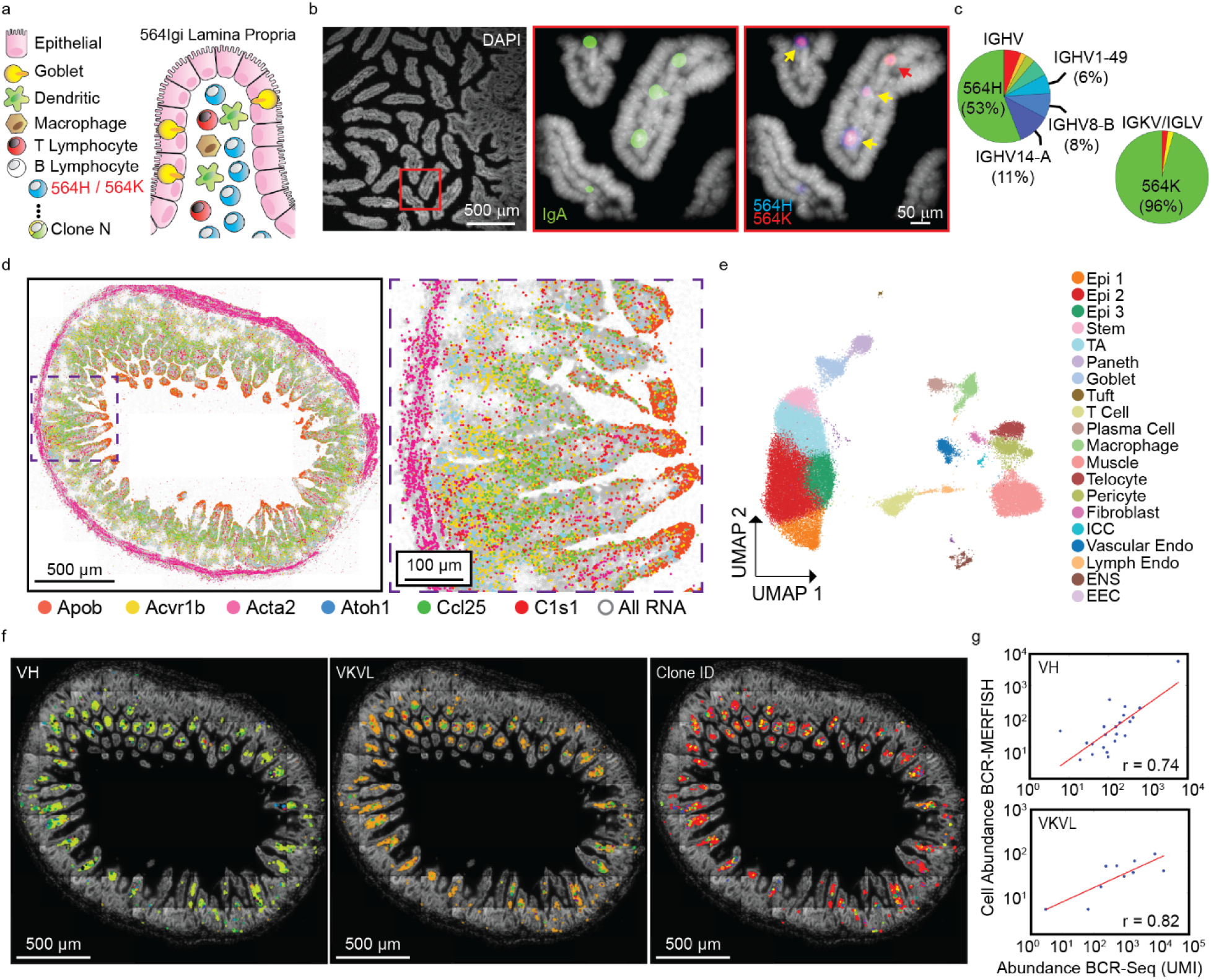
BCR-MERFISH accurately identifies V genes *in vivo*. **a**, Cartoon of a monogenic mouse model (564Igi) enriched in plasma cells expressing a prearranged BCR with a known VH and VK pair (564H/564K). **b**, Image of the ileum in a 564Igi mouse stained with DAPI (left) or in zoom-ins on the red boxed region with a probe marking IgA (green) and probes to the 564H (blue) and 564L chains (red). Yellow arrows mark 564H+/564L+ and the red arrow marks 564H+/564K-IgA+ plasma cells. Scale bars: 500 or 50 µm. **c**, Fraction of identified VH and VKVL genes in a 564Igi mouse assigned each V gene with BCH-MERFISH. **d**, Location of all mRNAs (gray) or 6 example mRNAs (color) identified with MERFISH in an ileal cross-section of an SPF mouse. Scale bars: 500 or 100 µm. **e**, UMAP of all cells profiled in three ileal slices of the mouse ileum colored by identified cell type. **f**, DAPI image (gray) of the ileal cross section with the boundaries of all identified plasma cells colored by VH (left), VKVL (middle), or the Clone ID (right) as determined by BCR-MERFISH. The V gene usage and abundance of all clones are listed in Table S5. **g**, The number of plasma cells assigned each VH (top) or VKVL (bottom) in three ileal cross sections with BCR-MERFISH versus the abundance of VH and VKVL genes observed in matched tissue determined from BCR-seq. r: Pearson correlation coefficients of the logarithmic abundances. Red lines: best linear fit.

We next sought to validate BCR-MERFISH in wild type mice by comparing plasma cell frequency determined with BCR-MERFISH to the frequency of V gene usage as determined via BCR-seq. Additionally, we designed V gene detection to be easily compatible with mRNA detection with standard MERFISH; thus, we also sought to validate our ability to combine these two modalities in BCR-MERFISH. To this end, we designed a 589-gene MERFISH library that includes canonical markers for all major cell populations in the mouse gut as well as near comprehensive coverage of mouse cytokines, chemokines, and their receptors (Tables S3&S4; Methods).

We then harvested the ileum from a wild type C57BL6 mouse harboring a native SPF microbiome and collected three slices for BCR-MERFISH and two intervening 100-µm-thick tissue blocks for BCR-seq (Methods). We performed BCR-MERFISH on these three ileal cross-sections, measuring successive barcode sets to identify mRNAs in all cells and the Fc region, VH gene, and VKVL gene in plasma cells (Fig. S1e-j). To identify cell boundaries, we included an antibody against a pan-cell surface marker—the Na^+^/K^+^-ATPase—and we leveraged these stains to define cell boundaries with Cellpose^43^ (Fig. S3a) and then refined this boundary-based cell assignments using the distribution and identity of the RNAs themselves with Baysor^44^ (Methods). Collectively, these measurements contained 55,291 cells.

We first examined the accuracy of mRNA measurements with MERFISH for samples co-stained with BCR-MERFISH probes. Classic marker RNAs were observed in their expected locations (Fig. 2d) and the abundance of RNAs correlated with bulk RNA-seq (Fig. S3b). Importantly, these measurements identified the cellular diversity expected for the ileum (Fig. 2e), and the identified cell clusters were marked by the expected genes (Fig. S3c). Thus, we conclude that the additional measurement of BCR-associated sequence features did not compromise our ability to accurately profile mRNAs and define cell types with MERFISH.

We next examined the properties of plasma cells determined via BCR-MERFISH in these slices. As expected for the ileum, we observed only IgA+ cells. Across all slices, BCR-MERFISH identified 23 VH gene groups and 33 VKVL gene groups, which collectively defined 98 plasma cell clones based on the unique pairing combinations (Fig. 2f; Fig. S3d; Table S5). Importantly, we observed reasonable clonal overlap between these slices (Fig. S3e), suggesting that the plasma cell diversity seen within the intervening blocks reserved for BCR-seq would be a representative reference for comparison.

We next performed BCR-seq on the paired tissue and compared the results from BCR-MERFISH (Methods). BCR-MERFISH detected a greater diversity of VH and VKVL gene groups than BCR-seq (Fig. S3f), perhaps reflecting the sequencing depth used. However, for the V genes detected with both methods, we observed a strong correlation between the number of plasma cells that use a specific VH or VKVL gene as detected via BCR-MERFISH with the frequency with which mRNAs associated with those genes were detected with BCR-seq (Fig. 2g). Thus, we conclude that BCR-MERFISH is capable of accurate identification of VH and VKVL gene groups *in situ*. Furthermore, as these measurements are made at a single-cell level, BCR-MERFISH naturally pairs heavy and light chain.

To confirm that BCR-MERFISH can identify plasma cell clones in other tissue types, we also performed measurements in mouse skin-draining lymph nodes (Fig. S4). Indeed, MERFISH was able to identify a diversity of cell types (Fig. S4a-d), identify the clonality of IgM+, IgA+, and IgG2+ plasma cells (Fig. S4e,f; Table S5), and use cell type distribution to define tissue anatomical features (e.g., light and dark zones, the subcapsular sinus, and the medullary cords; Fig. S4g). As expected, BCR-MERFISH revealed that the vast majority of plasma cells are found in the medullary cords region of the lymph node (Fig. S4e-h), indicating that BCR-MERFISH is capturing clones at the primary site of plasma cell expansion^24^. Interestingly, even within this small region, BCR-MERFISH revealed a diversity of plasma cell clones (Fig. S4h). Thus, we conclude that BCR-MERFISH is capable of simultaneous high-resolution gene expression measurements and plasma cell clone identification in a range of tissues.

### BCR-MERFISH maps microbiota-dependent plasma cell clonal diversity in the gut

To demonstrate the potential of BCR-MERFISH, we next sought to address some outstanding questions regarding the role that the microbiome plays in shaping plasma cell diversity in the gut. Plasma cells in the gut are primarily produced in gut-associated lymphatic tissues (GALT) such as PP and mesenteric lymph nodes (mLN)^45–47^. The presence of a gut microbiome is required for proper development of gut-associated lymphatic tissues (GALT) and the selection and affinity maturation of plasma cells, and the absence of a microbiome is linked to a marked reduction in plasma cell abundance within the small intestine^48^. However, the lamina propria of germ-free (GF) mice are not devoid of GALT, and plasma cell production continues to occur within PPs of GF mice^26,45^. However, it remains unclear to what degree the microbiome shapes the clonal diversity of plasma cells in the gut. In parallel, it has been shown that the PP and mLN GC of unrelated GF mice frequently include B cells with so called public clonotypes—common VH or VK genes, in some cases with identical junctional diversity and similar J usage^26,45^. However, it also remains unclear to what degree public clonotypes found in the GCs are also found in the plasma cell compartment of the gut, or how the presence or absence of a microbiome may shape the relative abundance of public clonotypes within the lamina propria.

To address these questions, we profiled ileal cross-sections from 2 SPF and 3 GF male mice (8wk, C57BL6) with BCR-MERFISH (Fig. 3a-d). As with the measurements above, we observed RNAs in the expected locations (Fig. 3a), strong correlation between replicates (Fig. S5a-d), and the expected cellular diversity across both conditions (Fig. 3b; Fig. S5e). Collectively, these measurements again support the ability to perform high-quality MERFISH in combination with BCR detection.

**Fig. 3.**
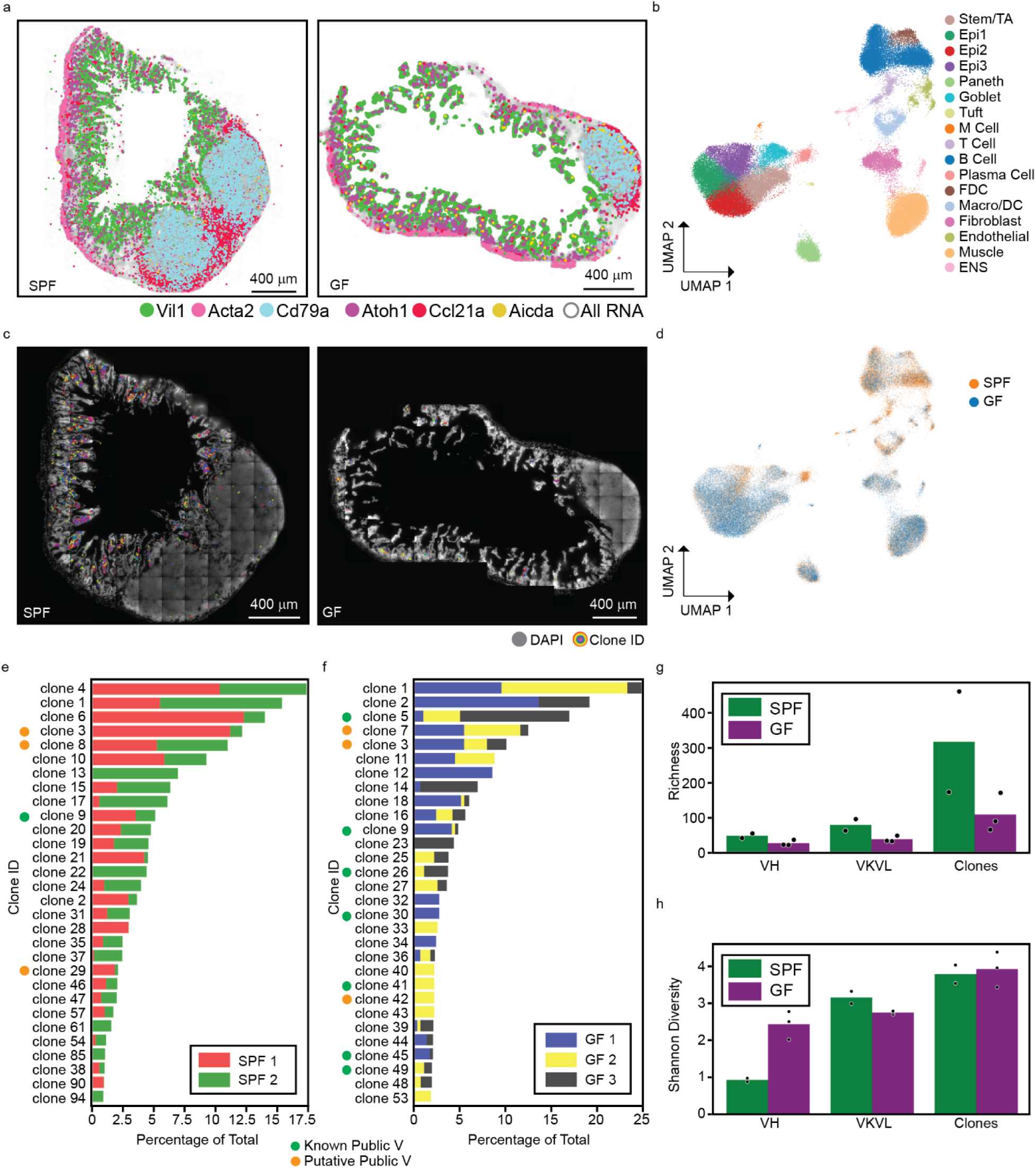
BCR-MERFISH reveals the microbiota-dependent modulation of plasma cell diversity in the gut. **a**, Location of all mRNAs (gray) or 6 example mRNAs (color) identified with MERFISH in an ileal cross-section of a SPF (right) or GF (left) mouse. Scale bars: 400 µm. **b**, UMAP representation of the cells imaged in all SPF and GF ileal sections colored by identified cell type. **c**, DAPI image of the slices in (a) with the clonal identity of plasma cells plotted in color. Scale bars: 400 µm. **d**, UMAP from (b) colored by microbiome composition. **e**,**f**, The abundance of each identified plasma clone across two SPF mice (e) and three GF mice (f). The V gene usage associated with each clone is in Table S5. Clones with unambiguous usage of previously reported public V genes are denoted with a green dot. Putative public clones identified by BCR-MERFISH are denoted with an orange dot. **g**,**h**, The average (bar) and individual (markers) richness (g) or Shannon Diversity Index (h) for VH, VKVL, and clonal IDs in all SPF and GF mice.

First, we explored the role of the microbiome on plasma cell diversity. As expected, BCR-MERFISH confirmed that the GF lamina propria is sparsely populated with IgA+ cells (Fig. 3c; Fig. S5f). Amongst the IgA+ cells, we decoded 540 and 271 different VH/VKVL pairs across all SPF and GF replicates, respectively (Fig. 3e,f; Table S5; Methods). Notably, some clones are shared between all mice. For instance, clone 1 (IGHV1-G/IGKV4-A) is within the top 5 most abundant clones for the SPF and GF conditions. By contrast, some clones were relatively abundant but were found only in a single SPF or GF mouse (clone 85: IGHV1-G/IGKV Collision143; clone 23: IGHV8-2/IGKV8-21, respectively).

To quantify the effect of the microbiome on clonality, we calculated the richness and Shannon Diversity Index for VH, VKVL and Clone IDs in SPF and GF mice (Methods). As expected, due to the large difference in plasma cell count (Fig. S5f), the total number of unique VH, VKVL, and Clone IDs (i.e., the richness) found in the SPF condition were all higher than the GF condition (Fig. 3g). However, despite a greater richness, the Shannon Diversity Indices found in SPF and GF mice were comparable for VKVL and Clone IDs (Fig. 3h). As Shannon Diversity is a measure of the relative abundance across the present clones, this observation suggests that there is a comparable degree of clonal expansion, albeit in a smaller number of unique clones, in GF as compared to SPF, which is consistent with the evidence for strong germinal center selection in PPs, even under germ-free conditions where GC activity is significantly damped^26^. Together, these results reveal that the microbiome drives a greater richness of plasma cell clones but plays a smaller role in the relative abundance of selected clones.

We next examined our measurements for evidence of the previously reported public clonotypes (Table S6). While our grouping of V genes may confound the identification of public VH or VKVL usage, we, nonetheless, found multiple plasma cell clones that used a public VH or VKVL associated with non-grouped genes in both SPF and GF mice (Fig. 3e,f; Methods; Tables S1, S5 & S6). For example, clone 9 (IGHV1-G/IGKV6-15) used a public VK gene and was found in both SPF and GF mice while clone 114, seen only in two GF mice, used the public IGKV6-15. Importantly the two GF public VH reported previously^26,45^ (IGHV1-12 and IGHV1-47) were seen in our GF measurements (clone 631: IGHV1-12/IGKV17-127; and clone 635: IGHV1-47/Collision 2). We also noted a marked enrichment of public clonotypes in GF mice. 5% (482 of 9645) of SPF plasma cells used a public V gene while 24.8% (157 of 633) of GF plasma cells did so. This observation is consistent with a similar enrichment observed in PP GC^26^.

Finally, our measurements also revealed a variety of non-grouped V genes that may represent new public clonotypes. We identified three putative public VK genes that were present across all SPF and GF mice: IGKV6-32, IGKV8-27, IGKV5-37 (Fig. 3e,f; Tables S5&S6). Notably, IGKV5-37 was paired with several different VH genes in the GF mouse (IGHV1-G, IGHV1-11, IGHV3-4, and IGHV8-2), suggesting that this VK gene segment was highly selected under GF conditions.

Collectively, these measurements provide one additional cross validation of the accuracy of BCR-MERFISH. Moreover, as the function of germinal centers in GALT is, in part, to create plasma cells that shape the secretory IgA landscape, these measurements now confirm that the emergence of public clonotypes in germinal centers does indeed shape the gut IgA repertoire. Notably, in exploring the usage of public clonotypes we observed that the vast majority of the previously reported public clonotypes were associated with non-grouped V genes and far fewer were associated with large V-gene groups, e.g. IGHV1-G, than would be expected via random chance (Table S6). This is important for two reasons. First, it suggests that there may be some selective pressure to use highly dissimilar V genes as public clonotypes, perhaps due to an existing propensity for binding to common or expected gut antigens. Second, it confirms that the grouping mechanism for BCR-MERFISH probe design minimally impeded our ability to identify public VH/VKVL genes. More broadly, these observations underscore the potential for BCR-MERFISH to discover trends in clonal expansion and diversity both in homeostatic and pathological conditions.

### Plasma cell clones are evenly distributed across the mucosal axis

A key advantage of BCR-MERFISH is its ability to precisely place plasma cell clones within specific tissue contexts and locations through its combination of gene expression profiling with MERFISH. Thus, to illustrate how MERFISH and BCR-MERFISH data can be jointly leveraged, we next explored the spatial distribution of plasma cell clones within precise regions of the small intestine. Antigen presentation to B cells is spatially organized within GALT structures, such as the PP or mLN, that receive antigens from different regions along the gastrointestinal tract^49,50^. However, plasma cells produced from these structures reach the gut indirectly via a route that involves migration through the lymphatics to the thoracic duct then recruitment from vascular circulation via the expression of the α_4_β_7_ integrin and the homing receptors Ccr9 and Ccr10^51^. Moreover, plasma cells can be found within different anatomical subregions of the gut, including multiple locations along the villi within the mucosa as well as the sub-mucosa. Thus, it is unclear whether the clonal identity of the secreted IgA is in any way organized within these anatomical features or along the length of the small intestine.

To explore clonal distributions along these axes, we harvested long strips of the ileum from two 8-week old SPF C57BL/6 males and characterized the distribution of plasma cell clones with BCR-MERFISH (Methods). To increase our mapping accuracy, we further filtered our IgA+ cells based on their cell-type label assignments as determined by clustering the single-cell MERFISH data (Methods). We identified genes in their expected locations (Fig. 4a), found that gene expression correlated strongly between the two replicates (Fig. S6a), and we were able again to define the expected cellular diversity in the gut (Fig. 4b; Fig. S6b,c), supporting our MERFISH measurements. In total, we imaged 301,911 cells of which 6,830 were plasma cells.

**Fig. 4.**
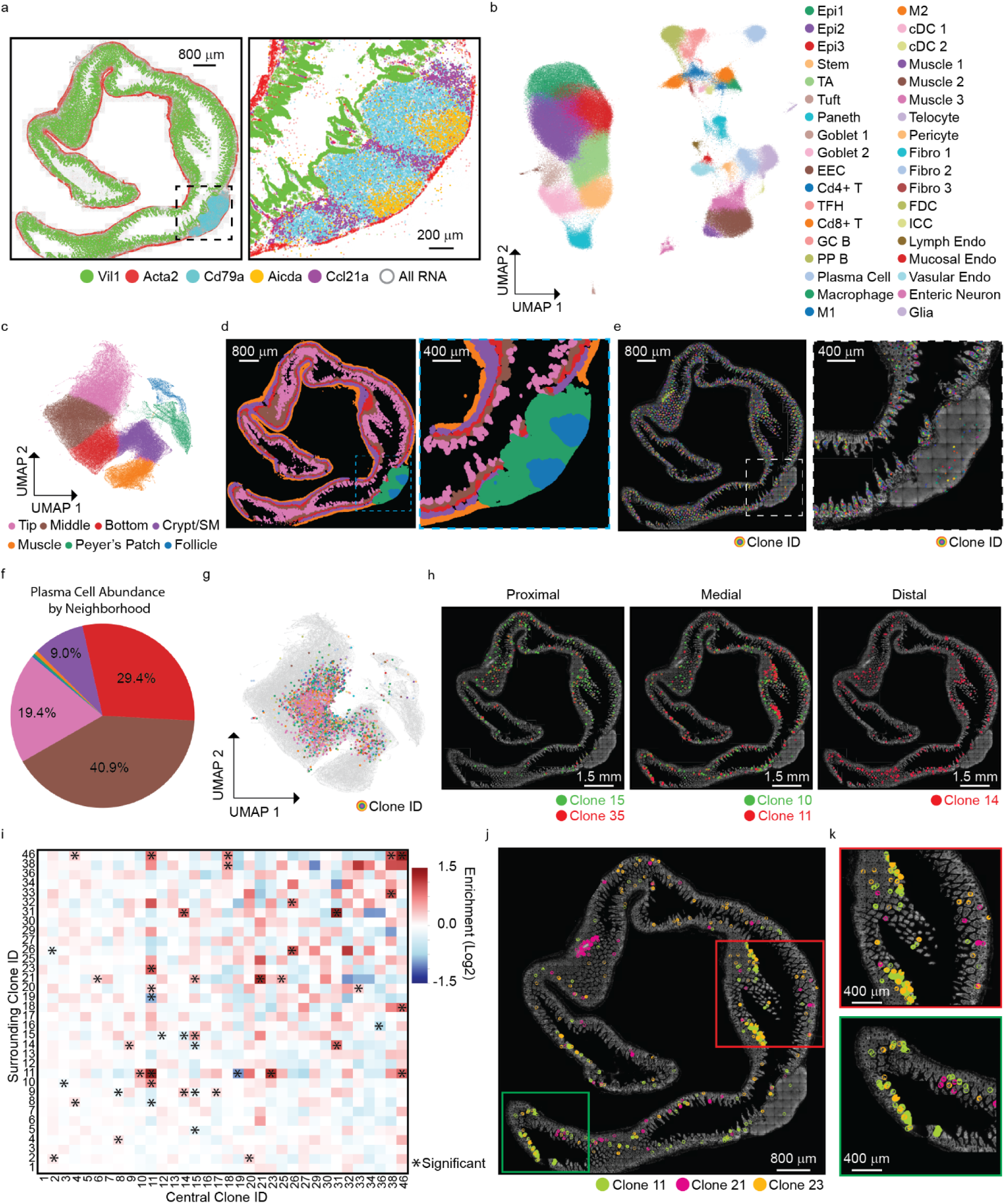
BCR-MERFISH reveals spatial heterogeneity in the distribution of plasma clones along the mouse ileum. **a**, Location of all mRNAs (gray) or 5 example mRNAs (color) identified with MERFISH in an ileal strip from an SPF mouse (right) with a zoom-in on a PP (right). Scale bars: 800 or 200 µm. **b**, UMAP of all cells profiled in two ileal strips colored by identified cell type. **c**, UMAP of neighborhood composition colored by the identified neighborhoods. **d**, Location of all cells in the ileal strip in (a) colored by neighborhood as in (c). Scale bars: 800 or 400 µm. **e**, DAPI images of the intestinal strips (gray) as in (a) with the location of all plasma cell clones (color) plotted. **f**, Fraction of plasma cells found in each neighborhood. **g**, Neighborhood UMAP as in (c) colored for all cells (gray) or plasma cell clones (colored by Clone ID). **h**, DAPI images (gray) of the slice in (a) with the location of clones found enriched in the proximal, medial, or distal ileum (colored by Clone ID). Scale bars: 1.5 mm. **i**, The enrichment in the spatial co-occurrence for plasma cells of one clonal type versus those of another for the strip in (a). Clone IDs are sorted by abundance in descending order. Columns represent the central clone while rows represent the clones in the surrounding neighborhood. **j**, DAPI image (gray) of the strip in (a) with the location of clones found to be spatially enriched next to themselves or other clones (color). Scale bar: 800 µm. **k**, As in (j) but for specific zoom-in regions.

To explore the organization of these plasma cells clones within different tissue regions, we leveraged a previously described tissue neighborhood detection algorithm that classifies tissue regions based on the local composition of cell types^36^. This analysis revealed all expected features in the gut defined by the expected cell populations (Fig. 4c-d; Fig. S7a-c). Namely, we identified neighborhoods associated with the top, middle, and bottom of the villus; a neighborhood comprising the crypts and sub-mucosa, and neighborhoods associated with the muscle layers, PP, and PP follicles (Fig. 4c-d; Fig. S7a-c).

Of the plasma cell identified via MERFISH, BCR-MERFISH revealed the clonal identity of 5,605 of them. Across both intestinal strips, we observed a diversity of plasma cell clones distributed across multiple tissue neighborhoods (Fig. 4e-g; Fig. S7d). Specifically, we identified 434 unique clone IDs collectively leveraging 47 different V gene groups and 45 different VKVL groups (Table S5). IGHV1-G dominated the VH usage, consistent with it being the largest group of VH genes whereas VKVL usage was much more uniformly distributed across V gene groups (Fig. S7e). As expected, the vast majority of plasma cells were found within the mid villus neighborhood with appreciable numbers also observed in both the top and bottom villi regions (Fig. 4f,g). This mid-villus enrichment likely reflects the spatial distribution of Ccl25 expression (Fig. 2d), which is the chemoattract associated with the homing receptor Ccr9 expressed on plasma cells^52,53^. In addition, we noted appreciable numbers of plasma cells in the Crypt/SM neighborhood, consistent with their entrance to the gut via the vasculature enriched in the sub-mucosa (Fig. 4f,g). We also noted a small number of IgA+ plasma cells outside and inside PP follicles (Fig. 4e-g). Whether these are resident or in transit remains to be explored.

However, plasma cell clones were well mixed between these regions, and we identified no statistically significant enrichment for any specific clone in any neighborhood (Fig. 4g; Fig. S7f; Table S7; Methods). On even shorter length scales, we also observed that the plasma cell clonality within individual villi was highly heterogeneous (Fig. 4e). Thus, we conclude the specific BCR carried by individual plasma cell does not play a role in their local tissue positioning in the healthy gut.

### Plasma cell clones are not uniformly distributed along the length of the ileum

We next explored whether there are variations in plasma cell clonal distributions along the length of the ileum. We first divided the ileal strips into evenly sized proximal, medial, and distal regions and explored the enrichment of individual plasma cell clones in these different regions for both strips (Fig. 4h; Fig. S8; Fig. S9). We noted modest variation in the richness and Shannon diversity of clones across these different regions (Fig. S8b-d; Fig. S9b-d). Nonetheless, we identified 57 Clone IDs that were statistically significant in their enrichment in specific regions of the ileum in either strip (Table S8). For instance, IGHV1-G/IGKV8-24 (clone 15) and IGHV1-G/IGKV1-88 (clone 35) were enriched in the proximal ileum, IGHV1-G/IGKV9-119 (clone 10) and IGHV1-G/IGKV8-21 (clone 11) were enriched in the medial ileum, and IGHV1-G/IGKV12-49 (clone 14) was enriched in the distal ileum in the first strip (Fig. 4h; Table S8, Methods). However, most clones were found distributed across the entire ileum (Table S8). Importantly, we observed regionally enriched clones in both strips (Fig. S9e; Table S8), indicating that a non-uniform distribution along the length of the ileum is likely a common property of SPF mice.

To explore whether any plasma cells may tend to co-occur in space, we computed the enrichment of different Clone IDs in the local neighborhood of each clone for one strip (Fig. 4i; Methods). Remarkably, this analysis revealed a variety of clones with specific pairwise spatial co-enrichment that was statistically significant (Table S9). For example, clone 11 (IGHV1-G/IGKV8-21) and clone 21 (IGHV1-G/IGKV Collision143) are statistically enriched next to themselves (Fig. 4i; Table S9) and were found in patches along the ileum (Fig. 4j,k). This analysis also found sets of different clones that co-occurred, suggesting a degree of higher-order organization (Fig. 4i-k). For example, clone 11 was also enriched next to clone 23 (IGHV1-G/IGKV Collision50), clone 14 (IGHV1-G/IGKV12-49) was enriched near clone 31 (IGHV1-G/IGKV3-7), and clone 18 (IGHV1-G/IGKV9-120) was enriched next to clone 46 (IGHV3-1/IGKV19-93). Importantly, similar patterns were seen, albeit with different clones, in the second strip (Fig. S10), confirming the generality of these distributions. As it is not expected that plasma cells will spatially co-occur or organize along the length of the ileum based on the targets of their secreted IgA, we hypothesize, instead, that this spatial patterning may be a reflection of the timing at which plasma cells enter the gut and, thus, spatial variation and correlation may reflect the temporally correlated selection and release of distinct clones from GALT. Nonetheless, while the mechanism of this patterning awaits further investigation, this observation underscores the discovery potential offered by BCR-MERFISH.

## Discussion

Here we introduced BCR-MERFISH, an imaging-based method built on top of the MERFISH platform for reporting the *in situ* clonality of plasma B cells by tracking the paired V gene usage within single cells. To achieve detection of highly homologous V genes, we developed a homology-aware probe design pipeline and a homology-resistant barcoding scheme. Importantly, the probe and barcode designs allow for simultaneous clonality tracking with standard MERFISH measurements for gene expression. We validated our V gene calling in cell culture, mice with a restricted BCR repertoire, and through paired BCR-sequencing of mice that express the native BCR repertoire. Furthermore, we showed that BCR-MERFISH can be applied to multiple tissues such as the ileum, PP, and skin-draining lymph node. We also used BCR-MERFISH to demonstrate that previously discovered public clones that appear within the germinal centers of unrelated germ-free mice also emerge in the plasma cell compartment of the germ-free lamina propria. Moreover, BCR-MERFISH identified additional putative public VK genes that emerge across SPF and GF plasma cells within the ileum. Finally, BCR-MERFISH uncovered an unexpected non-uniform distribution across relatively short stretches of the small intestine.

Many features of BCR-MERFISH complement those of the existing spatial clonality tracking tools. First, BCR-MERFISH provides a substantial extension in the number of B cell clones identifiably by previous microscopy-based methods, from the few clones identifiable with previous FISH methods^54^ to the 10 clones distinguishable via combinatorial genetic labels with Confetti mice^25,26^. Second, unlike Confetti, BCR-MERFISH defines clonality based on the sequence of the BCR itself. This feature may prove critical to link specific B cell clones identified via sequencing methods to the location of these cells within tissues. Moreover, as FISH probes can be designed against any RNA target; this feature should facilitate the application of BCR-MERFISH to a wide range of organisms, including those that are not easily genetically modified to express fluorescent determinants of clonality. Finally, BCR-MERFISH complements the recent spatial-capture based methods^27–30^ by providing single-cell resolution. This resolution is critical in providing an unambiguous linkage of heavy and light chain sequences in single B cells. Moreover, BCR-MERFISH is directly compatible with the high-resolution single-cell transcriptome measurements of MERFISH, and, thus, through combined measurements can directly determine the gene expression profiles of specific B cell clones as well as precise definitions of both the cellular and molecular niches that shape their behavior. For these reasons, we anticipate that BCR-MERFISH will prove to be a useful addition to the spatial-omics toolset for many tissue-level questions in adaptive immunity.

Nonetheless, BCR-MERFISH in its current form does have several important limitations. First, BCR-MERFISH provides a limited definition of BCR sequence. For example, we chose not to target D or J segments given their short length. As there are tens of D and J segments, their inclusion in future BCR-MERFISH variants would enhance the definable clonal diversity from thousands to millions. Future improvements to BCR-MERFISH may include these segments even with only a small number of probes by further leveraging the signal amplification provided by the high levels of BCR mRNAs in plasma cells. In addition, BCR-antigen affinity is set by nucleotide-level sequence features not revealed via V gene choice alone. FISH-based methods may not be the most suitable method for defining such sequence features, as the lack of *a priori* definition of these sequences and the relative insensitivity of hybridization to nucleotide-level variations will challenge probe design and targeting. Fortunately, as BCR-MERFISH defines B cell clonality via sequence features, it may be possible to couple sequencing-based methods—bulk, single-cell, or spatial—with BCR-MERFISH to link full sequence definition of individual B cell clones to their spatial location within tissue. Finally, BCR-MERFISH leverages the natural signal amplification produced by the high levels of plasma cell expression of BCR mRNA, yet the expression of BCR in naïve B cells or of TCR in T cells is much lower. However, as high-quality MERFISH has been performed for lowly expressed mRNA with only ∼10 probes^55^, we anticipate that our homology-aware probe design and encoding may be extended to enable the clonal definition of all B cells and perhaps T cells as well. Thus, we anticipate that BCR-MERFISH and its future extensions may provide new avenues for the study of role of adaptive immunity in contexts ranging from normal tissue homeostasis to cancer, infection, and autoimmunity.

## Acknowledgements

We thank members of the Moffitt and Carroll laboratories for helpful discussions. Portions of this research were conducted on the O2 High Performance Compute Cluster, supported by the Research Computing Group, at Harvard Medical School. This work was funded through National Institutes of Health grants (R01GM143277 to JRM; T32GM007753, T32GM144273, T32AI007529, and F30AI160909 to EA), a pilot grant from the Chan Zuckerberg Initiative to MCC and JRM, a postdoctoral mobility fellowship from the Swiss National Science Foundation to USH, and a grant from the Novartis Foundation for Medical-Biological Research to USH.

## Contributions

Conceptualization: EY, CC, USH, MCC, and JRM. Methodology: EY and JRM. Investigation: EY and JAS. Software: EY. Formal Analysis: EY. Validation: EY and JAS. Resources: EY, JAS, CC, USH, EHA, MCC, and JRM. Writing – original draft: EY and JRM. Writing – Review & Editing: EY, JAS, CC, USH, EHA, MCC, and JRM. Supervision: MCC and JRM. Funding acquisition: MCC and JRM.

## Methods

### BCR-MERFISH library design and construction

*In situ* B cell clonality was measured with separate sets of encoding probes that target the heavy chain and light chain variable (V) regions and encode the appropriate barcodes. Standard MERFISH encoding probes include a region of complementarity to the RNA target—the target region—and a barcode region comprised of a set of 20-nt readout sequences that define the barcode^31–34^. To address the high degrees of homology between V gene sequences, we developed a homology-aware target-region design algorithm which includes the following steps.

First, to create similarity trees for V genes, we downloaded the sequences for all IGHV, IGKV, and IGLV genes from the IGMT database^56^. We then constructed separate similarity trees for the IGHV genes or the IGKV and IGLV genes. We determined similarity between every pair of sequences by calculating the Jaccard index between the sets of all 13-mer subsequences that comprised each V gene sequence. We then defined a distance metric as 1 minus this similarity and leveraged agglomerative hierarchical clustering using the shortest distance method with a Euclidean distance metric to create the tree.

We then leveraged this similarity tree to guide usage of an existing target region design pipeline^32,33^ (github.com/ZhuangLab/MERFISH_analysis) to create target regions for V genes. Briefly, we used this pipeline to design 30-nt target regions with a GC content between 0.33 and 0.6 and a predicted melting temperature (TM) between 63 and 74 °C, allowing for target regions to overlap by as much as 20 nt. To penalize target regions for potential off target binding, we calculated a specificity or on-target score for each potential target region. This score was determined by considering two V gene sets: V genes within the same group (or node) as the V gene under consideration and all other V genes (the off-target group). For each possible 30-nt target region for a given V gene, we then calculated an off-target score defined as the ratio of the number of times each 13-mer subsequence within that possible target region was found in any of the off-target V genes normalized by the number of times each of these 13-mer subsequences were found in all V genes. We then calculated the specificity as 1 minus this off-target score. Conceptually, if a target region does not bind to any V gene in the off-target group its specificity is 1. Any potential binding to V genes in the off-target group (as judged by shared 13-mers) will decrease the specificity score towards 0.

We then used an iterative approach to group V genes based on the similarity tree such that each group of V genes would have at least ten target regions for every V gene within that group. As above, target regions were cut based on GC and TM. In addition, we cut target regions on specificity scores between 0.5 and 1. In the first iteration, we treated each V gene as its own group and designed target regions based on these cuts. For any V gene for which this approach did not generate 10 target regions, we identified the nearest neighbor on the similarity tree and grouped these genes. This process of target region design was repeated with the new specificity scores calculated to reflect these V gene groupings. Again, if after this iteration, any individual V gene did not have at least 10 possible target regions that satisfied the above constraints, it (or the group to which it belonged) was grouped with the nearest V gene (or potentially V gene group). This iterative process, which can be thought of as considering increasing large V gene groups in each iteration by stepping up the similarity tree one node per iteration, was run for a total of two times to identify groupings under which all VH and VKVL genes had at least 10 target regions. This produced 80 IGHV nodes and 68 IGKV/IGLV nodes (Table S1).

The barcodes assigned to the 80 IGHV nodes used a 14-bit, Hamming-Distance-4, Hamming-Weight-4 encoding scheme with 91 unique barcodes. 80 barcodes were randomly assigned to the IGHV nodes with the remaining barcodes assigned as ‘Blanks’ (Table S1). Similarly, the barcodes assigned to the 68 IGKV/IGLV nodes used a 14-bit, Hamming-Distance-4, Hamming-Weight-4 encoding scheme with 91 unique barcodes. 68 barcodes were randomly assigned to the IGHV nodes with the remaining barcodes assigned as ‘Blanks’ (Table S1). Individual readout sequences were associated with each of the 28 bits in these two 14-bit barcode sets, and encoding probes were designed by concatenating 3 of the 4 readout sequences associated with the barcode assigned to each V gene target to each of the target regions designed above. Finally, we concatenated unique priming sites to each encoding probe sequence such that we could amplify encoding probes from these templates (Table S2). The encoding probe templates were ordered from Twist Biosciences and amplified as described previously^57^.

### Encoding probes for constant regions

To augment plasma cell profiling, we designed an encoding probe set to capture Fc isotypes within plasma B cells. As there are only a small number of Fc regions, we encoded these regions with non-combinatorial barcodes. Specifically, we designed target regions against IgM, IgA, IgG1, IgG2c, IgG3, and IgE using the sequences for C57BL/6 mice from IMGT^56^. Target regions were designed against these sequences using the same target-region design pipeline discussed above with the same cuts on GC and TM. A single readout sequence associated with each Fc region was concatenated to these sequences, and these encoding probes were ordered synthesized directly from IDT as oligopools. IgD is associated with naïve B cells, where heavy chain expression is far lower and is detectable with combinatorial barcodes. Thus, we included this Fc in the combinatorial MERFISH library described below.

### Encoding probes for 564H and 564K

We designed single-molecule encoding FISH probes to target the germline sequences of the 564H and 564K V genes expressed by BCRs in the 564Igi transgenic mouse (Table S2). Target regions were designed using the same design pipeline discussed above, using the same parameters for GC and TM. Each probe was assigned a single readout sequence and probes were ordered synthesized directly from IDT as an oligopool.

### MERFISH library design and construction

We selected a panel of 589 genes to profile the gene expression patterns in the mouse small intestine, PP, and lymph node. This 589-gene panel included well-established cell-type markers for labeling major cell-type divisions in all tissues (e.g., epithelial, immune, muscular, neuronal, etc.) with additional genes selected to further discriminate between subdivisions of immune cell populations such as dark/light zone, naïve, and plasma B cells, as well as T cell and myeloid cell divisions. We also included all cytokines, chemokines, and their receptors, extracellular matrix receptors, cell adhesion molecules, pattern recognition receptors, and growth factors, with genes in the pathways largely derived from the KEGG database^58^. Since MERFISH data quality depends on RNA density within individual cells, we removed any RNA judged to be too abundant from companion RNA-sequencing (GEO: GSE143342)^59^. MERFISH encoding probes for this panel were designed using a previously described pipeline^26^. The target regions were 30-nt long, with an allowed GC content of 0.45-0.55, a predicted melting temperature between 65-75°C, and an allowed overlap between target regions of 20 nt.

To encode these genes, we used a 26-bit, Hamming-Distance-4, Hamming-Weight-4 encoding scheme with 640 unique barcodes. 589 of these barcodes were randomly assigned to genes while the remaining barcodes were assigned as ‘Blanks’ to allow for an internal measurement of the false positive rate (Table S3). We aimed to design 72 target regions per gene. Notably, some genes were not long enough to support 72 target regions, in which case, we included as many as possible. We assigned to each bit in these 26-bit barcodes a unique readout sequence, as described above (Table S2). However, to allow simultaneous profiling of these genes, VH, VKVL, and Fc regions, we ensured that these readout sequences were unique from those used above (Fig. S1e-j; Table S2). We randomly assigned 3 of the 4 readout sequences associated with each barcode assigned to a gene to each target region for that gene and concatenated priming sites to create template molecules for the encoding probes (Table S4). As above, these templates were synthesized by Twist Biosciences and amplified as described previously^57^.

### Tissue collection and cryosection

C57BL/6 mice were ordered as SPF or GF mice from Taconic Biosciences. SPF mice were housed in the Harvard Center for Comparative Medicine (HCCM) facility prior to sacrifice whereas GF mice were harvested upon receipt. 564IgI mice^42^ were bred and maintained by the Carroll laboratory and housed in the HCCM.

Mice were euthanized with isoflurane (Patterson, 07-890-8115) followed by cervical dislocation. Ileal tissue was harvested using a previously described fresh frozen protocol^57^. Briefly, the ileum (2 mm upstream of the cecum) was quickly harvested and flushed with 3 mL of 400 mM Ribonucleoside Vanadyl Complex (RVC; NEB, S1402S) in 1× phosphate buffered saline (PBS; Thermo, AM9625). The tissues were then placed in plastic cryomolds (Sakura, 4557), covered with Optimal Cutting Temperature Compound (OCT; Sakura, 25608-930), and immediately frozen with dry ice. After 30 minutes on dry ice, blocks were transferred to a –80 °C freezer.

Mouse lymph nodes were harvested differently. Following mouse sacrifice as described above, skin-draining lymph nodes were harvested and immersed in ice-cold 4% v/v paraformaldehyde (PFA; Electron Microscopy Sciences, 15714) in 1× PBS and incubated for 3 hours. The tissues were then incubated in 30% v/v sucrose (VWR, 0335-500G) and 4% v/v PFA in 1× PBS at 4°C for 12-16 hours. Tissues were then placed in plastic cryomolds, covered with OCT, and immediately frozen with dry ice. After 30 minutes on dry ice, tissue blocks were transferred to a – 80 °C freezer for storage.

10-micron-thick cryosections were generated with a Leica CM1850 cryostat prechilled to –20 °C. Slices were gently melted by touching the opposite side of the coverslip with a gloved finger and immediately refrozen by placing the coverslip on the cold surfaces of the cryostat chamber. As described previously^57^, slices were placed on 40-mm diameter coverslips (Bioptechs, 40-1313-03193) cleaned with hydrochloric acid and methanol and then coated with an allyl-silane layer (Sigma, 1077778), poly-lysine (Thermo, AM9625), and a layer of orange fluorescence beads (Thermo, F8800). The allyl-silane layer allows for covalent bonding of polyacrylamide films used for sample stabilization. The poly-lysine aids in tissue adherence, and the fluorescence beads serve as fiducial marks during MERFISH imaging. As an additional step for the PFA-fixed lymph nodes, the coverslips were dried for 30 minutes at room temperature.

Slices from both conditions were then fixed with 4% v/v PFA in 1× PBS at 4 °C for 10 minutes, then washed twice with 1× PBS. Tissues were permeabilized by incubating the coverslips in 70% ethanol at 4 °C for at least 12 hours. In some cases, samples were stored in 70% ethanol at 4 °C for up to one week before proceeding with subsequent steps. All water was RNase-free and was generated via reverse osmosis (Synergy UV, Millipore) with a Biopack polisher (Sigma, CDUFBI001).

All mouse experiments were performed in compliance with NIH guidelines and were reviewed and approved by the Harvard Institutional Animal Care and Use Committee (IACUC) under protocol IS00003215.

### MERFISH sample preparation

Following permeabilization for fresh frozen intestines and fixed lymph nodes, the 70% ethanol was aspirated and samples were washed twice with 30% v/v formamide (VWR, IC11FORMD002) in 2× Saline Sodium Citrate (SSC; Thermo, AM9765) at room temperature. The formamide buffer was then aspirated, and the coverslips were transferred into a parafilm-(Fisher, P7793)-coated Petri dish. An antibody and encoding library hybridization master mix was created and contains a 3 µg/mL of a monoclonal antibody against the Na^+^/K^+^-ATPase (Abcam, ab76020), 40 µM of the 589-gene MERFISH encoding probes, 1 µM of the IGHV encoding probes, 1 µM of the IGKV/IGLV encoding probes, 1 µM of the Fc encoding probes, and 2 µM of anchor probe in 2× SSC with 30% v/v formamide, 1 mg/mL yeast tRNA (Thermo, 15401029), and 10% w/v dextran sulfate (Sigma, S4030). The anchor probe was the same as that used previously^57,60^ and had a sequence of/5Acryd/TTG AGT GGA TGG AGT GTA ATT+ TT+ TT+ TT+ TT+ TT+ TT+ TT+ TT+ TT+ T, where /5Acryd/ represents an acrydite modification and T+ indicates locked nucleic acid (IDT). 60 µL of this hybridization master mix was added to each coverslip. To avoid evaporation of the hybridization solution, a wet Kimwipe was inserted inside the Petri dish. The sample was hybridized for 24 hours at 37 °C in a humidified oven. Unbound probes were removed through two washes in 30% v/v formamide in 2× SSC, each for 30 minutes at 47 °C.

For immunofluorescent labeling of cell boundaries, we generated a methacrylate-modified secondary antibody inspired by previous methods for preserving antibody signal in samples cleared via proteolysis^61^. Methacrylate NHS (Sigma, 730300-1G) was diluted to 100 mM in anhydrous DMSO (Thermo, D12345), and a goat anti-rabbit IgG labeled with Alexa488 (Thermo, A32731) was mixed with 100 mM MA-NHS at a 2:1 volumetric ratio and incubated in the dark for 1 hour at room temperature. This labeled antibody was then diluted 10-fold into 2× SSC with 30% v/v formamide, 1 mg/mL yeast tRNA (Thermo, 15401029), and 10% w/v dextran sulfate (Sigma, S4030), and samples were incubated in this solution for 24 hours at 37 °C in a humidified oven. Samples were then washed in 2× SSC at room temperature.

Samples were stabilized and cleared as described previously^32,57^. Briefly, they were washed twice with a hydrogel solution consisting of 4% 19:1 acrylamide/bis-acrylamide (Bio-Rad, 1610144) in 50 mM Tris-HCl (Thermo, 15568-025), 300 mM NaCl (Thermo, AM9759), 0.03% w/v ammonium persulfate (Sigma, 215589), and 0.15% v/v tetramethylethylenediamine (TEMED; Sigma, T7024). The coverslips were then inverted onto a 65 µL droplet of this hydrogel solution spotted onto a GelSlick-(Lonza, 50640)-coated glass plate (Gorilla Scientific, 6101). This gel solution was covered from light with aluminum foil and allowed to polymerize for 2 hours at room temperature. Coverslips were removed from the glass plate using a razor and tweezers and placed inside a 60 mm Petri dish containing a digestion buffer comprised of 1:100 proteinase K (NEB, P8107S) in 0.25% v/v Triton-X (Sigma, T8787), 2% v/v Sodium Dodecyl Sulfate (SDS; ThermoFisher Scientific, AM9823), and 2× SSC. Samples were incubated at 37 °C for two days. Digestion buffer was refreshed every 24 hours. After digestion, the sample was washed with 2× SSC for 30 minutes at room temperature for a total of 6 washes. Extensive washing is needed to remove residual SDS. After the final wash, the samples were stored in 2× SSC at 4°C. Samples were stored for no longer than 1 week prior to imaging.

### MERFISH imaging

To prepare samples for imaging, the 2× SSC was aspirated from each sample and replaced with 5 mL of readout hybridization buffer consisting of 10% v/v ethylene carbonate (Alfa Aesar, A15735-36) in 2× SSC, 0.25% v/v Triton-X (Sigma, T8787), and 3 nM of each of the fluorescently labeled readout probes associated with the first two bits used in the 589 gene MERFISH library. Samples were stained for 30 minutes in the dark at room temperature with rocking. The staining solution was aspirated and washed for 15 minutes in a readout buffer comprising 10% v/v ethylene carbonate and 0.25% v/v Triton-X in 2× SSC.

The sample was then stained with 1 mg/mL DAPI (FisherScientific, D1306) for 10 minutes with rocking, washed once more with 2× SSC and loaded into the MERFISH system. To prepare the MERFISH fluidics device, individual wells of a 24-deep-well plate (Westnet, 504361) were filled with 6 mL of the readout hybridization buffer described above with each well containing a different pair of readout probes associated with the bits of the 589 gene panel. The sequences of these probes have been described elsewhere^32,34^ and are reproduced in Table S2. Readout probes were synthesized by Biosynthesis, Inc and the indicated fluorophores were linked to the oligonucleotides via a disulfide bond to allow their cleavage and removal between imaging rounds. The 24-well ‘readout cartridge’ was loaded into a custom manifold and sipper system described previously^36,62^.

Additionally, four more buffers were created, placed into 50-mL falcon tubes, and loaded into the fluidics system. The first buffer is a TCEP-based cleavage buffer to cleave the disulfide bond that anchors fluorophores to readout probes, effectively removing the source of the fluorescence for each bit between imaging rounds. This buffer comprised 50mM tris(2-carboxyethyl)phosphine (GoldBio, TCEP25) in 2× SSC. The second is an imaging buffer comprised of 4 mM Trolox-quinone, 0.5 mg ml/mL Trolox (Abcam, AB120747), 1:500 recombinant protocatechuate 3,4-dioxygenase (rPCO; OYC Americas, 46852004) and 5 mM protocatechuic acid (Sigma, 37580-25G-F) in 2× SSC. Trolox-quinone was generated via UV incubation of a solution of Trolox as described by published approaches^57,63^. The final two buffers were the readout wash buffer described above and 2× SSC, which was used to wash away excess cleavage buffer.

These samples were imaged on a microscope system described previously^36,57^. The desired fields of view (FOVs) were selected from a tiled mosaic created of the sample using the DAPI channel, 405-nm illumination, and a 10× objective. All selected FOV were then imaged with a 60× objective (Nikon, 1.4NA oil ApoChromat) to collect a z-stack in each image channel for each FOV. The z-stack comprised seven to ten 1.5-mm-spaced z-planes. Samples were imaged on a microscope that leveraged two cameras split with a Cairn TwinCam and the Alexa750 and Cy5 channels were imaged simultaneously with the system using the 635-nm and 735-nm illumination of a Celesta light engine (Lumencor). Similar z-stacks were collected in the 405-nm and 473-nm channels in the first round of imaging to image the nuclei with DAPI and the Na^+^/K^+^-ATPase with the NHS- modified Alexa488 secondary antibody. An image of the fiducial beads was also collected with the 535-nm illumination to reregister all images across all imaging rounds. After all FOVs were imaged, an automated flow system introduced the cleavage buffer and incubated the sample for 20 minutes, washed the sample with 2× SSC, introduced the next readout hybridization buffer and incubated the sample for 30 minutes, and then washed the sample with readout wash buffer for 6.5 minutes. Imaging buffer was then introduced and the process repeated, imaging just the 750-nm, 635-nm, and 535-nm channels. This process was used to first image the 26-bits associated with the 589-gene panel. All buffers were then refreshed and a new readout reagent cartridge loaded into the system. This process then continued to collect the 14 channels required for the VH barcodes, the 14 channels for the VKVL, and the 6 channels required for the Fc regions.

### V gene detection in cell culture

To validate our ability to identify V genes using BCR-MERFISH, we created a plasmid set that encode prearranged heavy and light chains under constitutive expression of the CMV protomer. To create these plasmids, we downloaded the IGHV, IGKV, and IGLV genes for C57BL/6 mice from IMGT^56^. We then ordered a cloning template for each V gene using these sequences as individual gene blocks from Twist Biosciences. We used Gibson assembly to insert each of these V genes into the pSeq8 plasmid, which contains a fully assembled heavy chain gene with the IgG2 constant region, the ColE1 origin of replication, and a kanamycin resistance cassette.

We performed this cloning using a pooled approach, transformed this plasmid pool into *E. coli*, and selected colonies on LB + kanamycin plates. Plates were sent to Genewiz for colony picking, plasmid purification, and sanger sequencing of the V gene inserts. Individual clones were selected based on these sequencing results, and plasmid preps of these bacterial strains were created by Genewiz. Notably, >90% of the IGHV, IGHK, and IGLV segments were successfully cloned into the pSeq8 backbone in this pooled cloning reaction.

To then complete this set, we identified V gene segments not incorporated into pSeq8 and used the same Gibson assembly and selection approach but with individual V genes. The result was a set of expression plasmids each of which expressed a prearranged heavy chain with one of the 188 IGHV and 102 IGKV/IGLV regions. We created glycerol stocks for each plasmid by transforming competent *E. coli* cells (Thermo Fisher, C737303).

We then selected defined subsets of these plasmids, prepared them from *E. coli* stocks using standard mini preparation protocols (Zymo, D4015), and transfected individual plasmids into HEK293 cells. Briefly, we seeded individual wells of a 24-well culture plate (Corning, 353047) with 25,000 cells per well in 1 mL of a culture media comprised of DMEM (ATCC, 30-2002) supplemented with 10% fetal bovine serum (FBS; Thermo, MT35010CV) and 4 mM L-glutamine (Thermo, 25030149). Cells were cultured at 37 °C for 24 hours.

We then created a transfection mixture for each of the desired plasmids using 100 µL of OptiMEM (Thermo, 31985062), 500 ng of each plasmid, and 1 µg of polyethylenimine (R&D Systems, 7854/100). Each mixture was incubated in the dark for 30 minutes before being added dropwise to the appropriate culture well. Each plate also contained a single well that was transfected with a pCMV::GFP plasmid to serve as a transfection control (Addgene, 11153). Cells were returned to the 37 °C incubator for 24 hours. The following day, the media was aspirated from each well and replaced with 50 µL of Trypsin (Thermo, 25-200-114), and the plates were returned to the 37°C incubator for 5 minutes.

The de-adherence of the cells from the bottom of the plates was then monitored with light microscopy, and when the cells were no longer adherent, 1 mL of culture medium was added to each well. The media was then collected from each well, pooled into a single 50 mL falcon tube, and the cells were concentrated by spinning at 1000×g for 2 minutes. The media aspirated and the cells were resuspended in culture media to a final concentration of 100,000 cells/mL. 5 mL of this cell mixture was then placed on a 40-mm coverslip which was silanized and treated with poly-lysine, as above, in a 60 mm petri dish. Cells were allowed to adhere and grow on this coverslip at 37°C for 24 hours.

To prepare cells for imaging, **t**he media was aspirated, the coverslips were washed 3 times in 1× PBS, and the samples were fixed with 3 mL of 4% v/v PFA in 1× PBS at room temperature for 10 minutes. Fixation solution was then removed with three washes with 1× PBS, and the cells were permeabilized in a mixture of 0.5% Triton-X in 1× PBS at room temperature for 15 minutes. Staining and imaging of the samples was performed using the protocols described for tissue samples; however, no antibodies were used in the preparation of these samples.

### Decoding of individual RNAs

RNAs included in the 589-gene MERFISH panel were decoded from these raw images using a previously described pipeline (github.com/ZhuangLab/MERFISH_analysis)^32,33,57^. Briefly, the fiducial bead images were used to generate affine transformations to correct small stage offset between imaging rounds in the same FOV. These transforms were used to create a tiff stack for each FOV in which all imaging channels associated with the barcodes for all V genes, the Fc regions, and the 589-gene MERFISH panel as well as the DAPI and Na^+^/K^+^-ATPase co-stain images were warped to a common coordinate system. The image channels associated with the 589-gene MERFISH panel were then normalized to correct for intensity variations between rounds and the intensity across these images for each pixel was matched to that predicted for the possible barcodes to assign pixels to barcodes. Background pixels were removed based on their Euclidean distance to the nearest barcode and adjacent pixels assigned to the same barcode were aggregated to form putative molecules. Background molecules were excluded by cutting on the number of pixels assigned to individual molecules and the average brightness across these pixels, compiled across all imaging rounds via the Euclidean norm. These thresholds were adjusted per dataset.

### Segmentation of tissue images

Cell boundaries were identified in tissues in two steps. First, we used Cellpose v2.2.1^43,64^ to identify cell boundaries on the Na^+^/K^+^-ATPase immunofluorescent images. Specifically, we started with the cyto3 model and then used a human-in-the-loop training model in which we corrected cellpose segmentation results for a set of FOVs taken from PP and the surrounding ileum^64^. The model was trained for 5000 epochs and we used the following parameters: centerZ=5, minZ=1, maxZ=10, inputDiameter=68. Cellpose was run on individual z-planes, and then we used the overlap between cellpose masks between z-planes to stitch together 3D boundaries. RNAs were assigned to these 3D masks if they fell within their boundaries.

We then refined these cell boundaries using the identity and spatial distribution of the RNAs identified in the 589-gene MERFISH panel using Baysor v0.4.3^44^. Baysor was run on individual datasets with the following parameters: scale=4, scale STD=100%, nclusters=10 and a prior confidence associated with the RNA-cell assignment generated with the cellpose masks of 0.9. As Baysor will adjust the mRNAs associated within individual cells relative to the assignment made via cellpose and will be able to assign mRNAs to cells that were not identified with Cellpose, we leveraged the Baysor cell assignments to generate final masks for individual cells. Specifically, we created alpha hulls using the python package Alphashape with shape parameter alpha=0.3.

### V gene decoding

To decode the barcodes associated with V genes, we developed an approach that leveraged the very high expression level of BCR transcripts in plasma cells. First, we integrated the average fluorescence in each alphahull for each cell across the Fc images to generate an observed optical barcode. We used a fixed threshold of 15% maximum pixel intensity across any of the channels to determine Fc+ plasma cells. For these cells, we then integrated the average fluorescence intensity within these boundaries across each of the imaging rounds associated with the two V gene barcode sets. For each V gene barcode set, we then calculated the normalized fluorescence intensity using the Euclidean norm. For each cell, we then computed an estimated HW by thresholding on these normalized intensities using a fixed threshold of 35% maximum pixel intensity. Cells that had observed optical barcodes with an apparent HW greater than 4 were labeled as ‘Collision’ cells and set aside. Rare cells with observed optical barcodes with an HW of 2 or less based on this threshold were not analyzed further. All other cells were then assigned to a V gene group by selecting the nearest V gene barcode to the observed optical barcode using a Euclidean distance.

To then discriminate among the ‘Collision’ cells, we created a comprehensive list of all possible pairwise combinations of valid V gene barcodes, i.e., possible collision barcodes (Table S1). As above, we then matched each of the normalized optical barcodes for all ‘Collision’ cells to the expected normalized optical barcodes associated with all pairs of valid V gene barcodes to assign each cell a unique collision barcode label. All alphahulls were then assigned a Clone ID based on its co-expression of decoded VH and VKVL barcodes.

For the neighborhood and spatial enrichment analysis for the intestinal strips, we extended the decoding pipeline to integrate the cell-type assignments determined by MERFISH. Specifically, we adopted a more stringent definition of plasma cells to avoid potential spatial co-variation artifacts that might arise when portions of the intense Fc, VH, or VKVL staining might spill into to adjacent non-plasma cells. First, we labeled all cell-types using the RNA as determined by MERFISH. Then, we extracted the alpha hulls for all Fc+ cells, as above, subset them based on those that had sufficient staining in both the VH and VKVL channels and determined their Clone IDs through their V gene co-expression. Finally, we kept only the cells that were assigned to the Plasma Cell cluster as determined via MERFISH.

For the 564Igi mice, we modified the above pipeline slightly to identify V gene usage in these measurements. Specifically, the plasma cell density within the lamina propria of 564Igi mice was sparse enough that we could segment plasma cells using the intensity of the Fc stain directly with a thresholding mask based on Otsu’s method. Once boundaries were identified the VH and VKVL optical barcodes were assembled and identified as described above. The same approach was used to detect V genes in the transfected HEK293 cells.

### Single-cell analysis

Cells were identified using standard single-cell approaches provided with the scanpy pipeline^65^. Briefly, gene expression profiles for the 589-gene MERFISH library were used for all cell identification. A gene expression matrix was computed by adding the abundance of all mRNAs within individual cells. Cells were then filtered to keep those that expressed more than 28 or less than 650 mRNAs. Gene expression was then normalized by the total number of mRNAs measured per cell and scaled by proportion (dividing each cell’s gene count by the total count for that cell). For all datasets, we combined cells from multiple replicates without the application of batch correction. Expression was then log normalized after the addition of a single pseudo-count, and expression was z-scored. A principal component analysis (PCA) was then performed to reduce the dimensionality of this expression matrix, and we kept a variable number of principle components selected to optimize properties of the UMAP visualization for each dataset. We calculated a UMAP using standard parameters in scanpy using the remaining principal components (PC), and then applied Leiden clustering on each dataset with a resolution hand-selected to optimize the division of the expected cell types. Cell types were labeled based on the expression of marker genes, which were calculated using standard scanpy methods.

To identify the cellular neighborhoods within different tissues, we leveraged a neighborhood identification method introduced previously^36^. Briefly, we computed a neighborhood composition vector defined by the abundance of all cell types within the 80 nearest neighbors of each cell. These vectors were then decomposed with a PCA, 43 PCs were kept, and a UMAP visualization and Leiden clustering were performed as above. Neighborhoods were labeled based on the location in the tissue and their cell type composition.

V gene optical barcode UMAPs were calculated using the normalized intensity observed for individual cells for either the VH or VKVL images. These values were decomposed via PCA, all PCs were kept, a nearest neighbor graph was constructed, and a UMAP constructed using the standard scanpy parameters as described above.

### BCR-sequencing

An ileal block from an SPF mouse was prepared as described above. To create paired BCR-MERFISH and BCR-sequencing datasets, we collected tissue samples for each measurement as follows: during cryosection, one 10 µm slice was placed on a coverslip to be imaged with BCR-MERFISH, then the immediate next 100 microns of tissue (10 slices) was placed in a tube containing TRIzol (Thermo, 15596026) for BCR-sequencing, and then the next slice was collected and placed on the same coverslip for BCR-MERFISH, then the next 100 microns were placed in the TRIzol tube, and then finally a third slice was placed on the coverslip for MERFISH. In total, we collected 200 microns of tissue for BCR-sequencing, which was flanked by three slices for BCR-MERFISH. Tissue collected for BCR-sequencing was immediately immersed in Trizol and total RNA was extracted via column purification (Zymo, R2050). To prepare samples for BCR-sequencing we used a commercial BCR-sequencing preparation kit (NEB, E6330S) with.cDNA constructed following the manufacturer’s instructions. cDNA was then sequenced by the Biopolymers Facilities at Harvard Medical School on the MiSeq Illumina instrument with 600 nucleotide-long paired-end reads.

To analyze these BCR-sequencing results, we used the pRESTO^66^ pipeline. Briefly, we aligned reads to V genes in the C57BL/6 mouse using IgBLAST^67^, and then extracted the Fc for heavy chain reads. We subset our heavy chain reads to only those associated with IgA, as we only observed IgA positive plasma cells within the lamina propria. For light chains, we performed similar analysis, except we did not filter on IgA+ reads.

To then compare the abundance of V genes measured with BCR-seq to the frequency with which we observed plasma cells containing specific light and heavy V gene nodes with BCR-MERFISH, we compiled the total abundance associated with each heavy and light V gene determined BCR-seq, then summed the total abundance associated with all V genes assigned to BCR-MERFISH groups (Table S1).

### Calculation of plasma cell richness and Shannon Diversity Index

To determine the richness and Shannon diversity associated with plasma cell clonality in SPF and GF ileal cross sections, we performed the following calculations for each replicate individually. The richness was calculated directly from the number of unique clone IDs in each dataset. The Shannon Diversity Index (SDI) was calculated using the following equation,

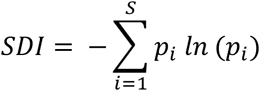

where *p*_i_ is the proportion (fraction) of the *i*th clone in the population of all *S* clone IDs observed and *ln* represents the natural logarithm.

### Regional plasma cell enrichment

To determine if any plasma cell clones showed a regional enrichment along the length of the ileum, we separated each of the two ileal strips into proximal, medial, and distal regions using manual cuts on location, selected such that the three regions had comparable areas. Richness and the Shannon Diversity Index were calculated for each region in each of the two replication measurements as described above. To determine if any given clone ID was enriched in the proximal, medial or distal ileum, we used a chi-squared test, and we corrected for multiple hypothesis testing using Benjamini-Hochberg corrections. We used a significance threshold after multiple hypothesis correction of p > 0.05.

### Spatial co-occurrence of plasma cells

To determine if any pair of plasma clones tended to co-occur in the same local regions of the ileum, we calculated a cell-to-cell proximity enrichment for each pair of clone IDs. For each plasma cell, we identified the clone ID of the nearest 100 plasma cells to determine the number of each clone found within its neighborhood. To compute the average neighborhood composition for each clone, we then averaged the neighborhoods for all clones of the same ID. To then determine an enrichment, we divided these clonal neighborhoods by the average abundance of all clone IDs.

To determine the statistical significance of these enrichment values, we used a randomized resampling method. Specifically, we repeated the calculation above 10,000 times. In each of these 10,000 resamplings, we maintained the location of plasma cells but randomly reassigned all identified clone IDs to different plasma cells. We used these resamplings to estimate the null distribution for enrichment between each pair of clone IDs, and then used this null distribution with a two-tailed t-test to calculate the significance for the measured enrichment between a given clone pair calculated with the real clone IDs assigned to each plasma cells. We corrected these p-values with the Benjamini-Hochberg correction and used a significance threshold of 0.05, as described above.

