## Supplementary Information and Figures for "In situ profiling of plasma cell clonality with image-based single-cell transcriptomics"

### Supplementary Figures and Captions

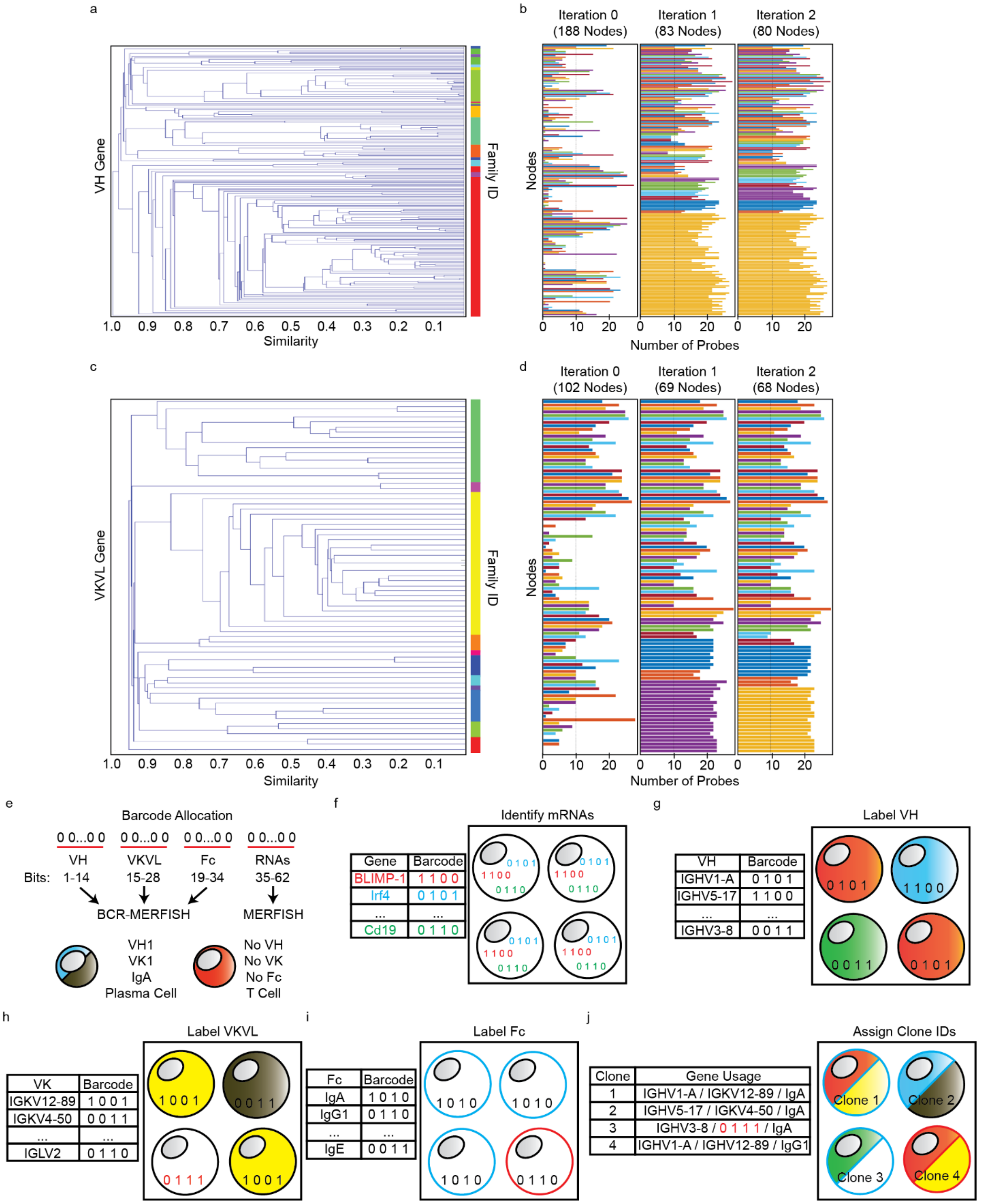

**Fig. S1 | Design of VH and VKVL target regions and the BCR-MERFISH strategy. a**, Similarity tree defining the relative sequence homology between mouse VH genes (left). The family assignment for each V gene is indicated by color (right). **b**, The number of target regions that can be designed against each VH gene if all VH genes are treated independently (left), after one round of grouping VH genes with the nearest neighbor on the similarity tree if they cannot support sufficient target regions (middle), and after an additional iteration of this process (right). The dashed line represents the arbitrary threshold of 10 target regions per gene used in this work. Color indicates the group to which the V gene has been assigned. V genes are ordered as in (a). **c**, Similarity tree as in (a) but for VKVL genes. **d**, The number of target regions versus iterations as in (b) but for VKVL genes. **e-j**, Schematic depiction of how BCR-MERFISH stacks multiple barcode sets (e) to allow the identification of hundreds or thousands of individual RNA molecules across all cells in a tissue via standard MERFISH (f) and, within plasma cells, the identification of the V gene choice for the heavy (g) or light (h) chain, and the identification of the specific Fc choice for the heavy chain (i). Collectively, the combination of gene expression profiles as determined via MERFISH (f) and BCR features as determined here (g, h, and i) allow the mapping of B cell clones in the context of complex tissues (j). The red barcode in (h) illustrates a situation in which the homology-resistant barcoding strategy detected a collision. This collision is marked by a Hamming Weight greater than the expected HW of 2 used strictly for this example cartoon. There are many unique collisions (Table S1) so individual collisions can still be used to define B cell clonality.

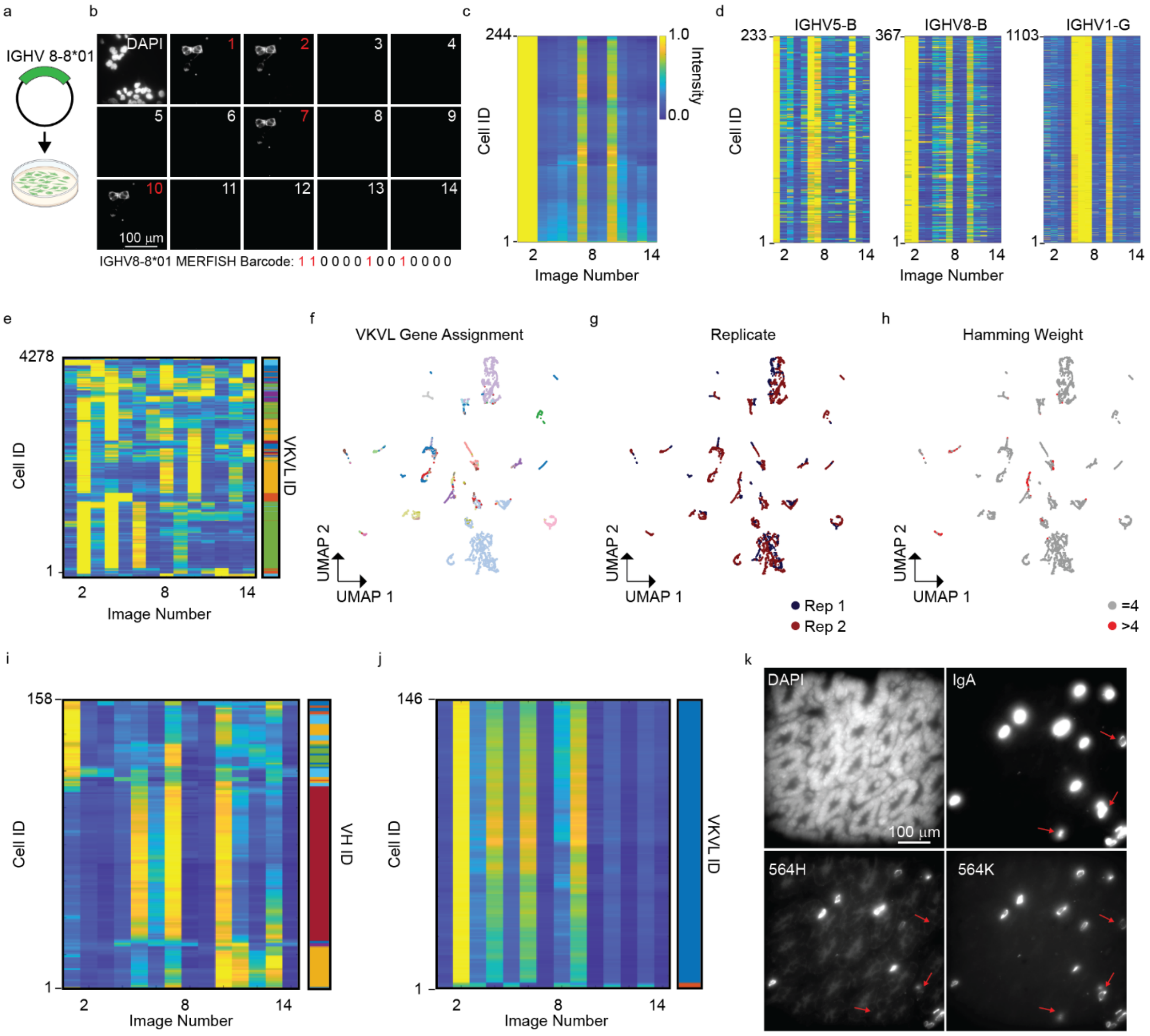


**Fig. S2 | Validation of V gene identification in cell culture and in mice. a,** Illustration of the transfection of a single plasmid expressing a prearranged heavy chain using IGHV8-8*01 into HEK293 cells. **b**, The fourteen images used to decode the VH gene in HEK293 transfected with the plasmid in (a). A DAPI image highlights the location of all cells, and red highlights the images associated with the expected barcode, which is listed on the bottom. Scale bar: 100 µm. **c**, Normalized intensity for each transfected cell across all images associated with the identification of VH genes for the measurements in (a,b). **d**, Normalized intensity across all images used to identify VH genes for all transfected cells in a mixed plasmid experiment (Fig. 1f) assigned to the listed V gene groups in the titles. The intensity scale is as in (c). **e**, The normalized intensity across all images used to identify VKVL genes (left) for all HEK293 cells transfected with a mixture of plasmids expressing a prearranged heavy chain that uses defined VKVL genes with the VKVL ID assigned via BCR-MERFISH (color, right). **f-h**, UMAP representation of the normalized intensity profiles in (e) colored by VKVL gene assignment (f), replicate ID (g), and the measured HW of the optical barcode (h). **i**,**j**, The normalized intensity across all images used to assign VH (i) or VKVL (j) for IgA+ plasma cells measured in the ileum of 564Igi mice (left) or the V gene assigned to them based on their optical barcode (color, right). **k**, Representative images of an ileal region of 564Igi mice stained with DAPI (top left), Fc probes targeting IgA (top right), probes targeting the expected VH gene for 564Igi mice (564H, left bottom), and probes targeting the expected VK gene for 564Igi mice (564K, right bottom). Arrows mark IgA+ cells that express the 564K gene but not the 564H gene, as seen with BCR-MERFISH. Scale bar: 100 µm.

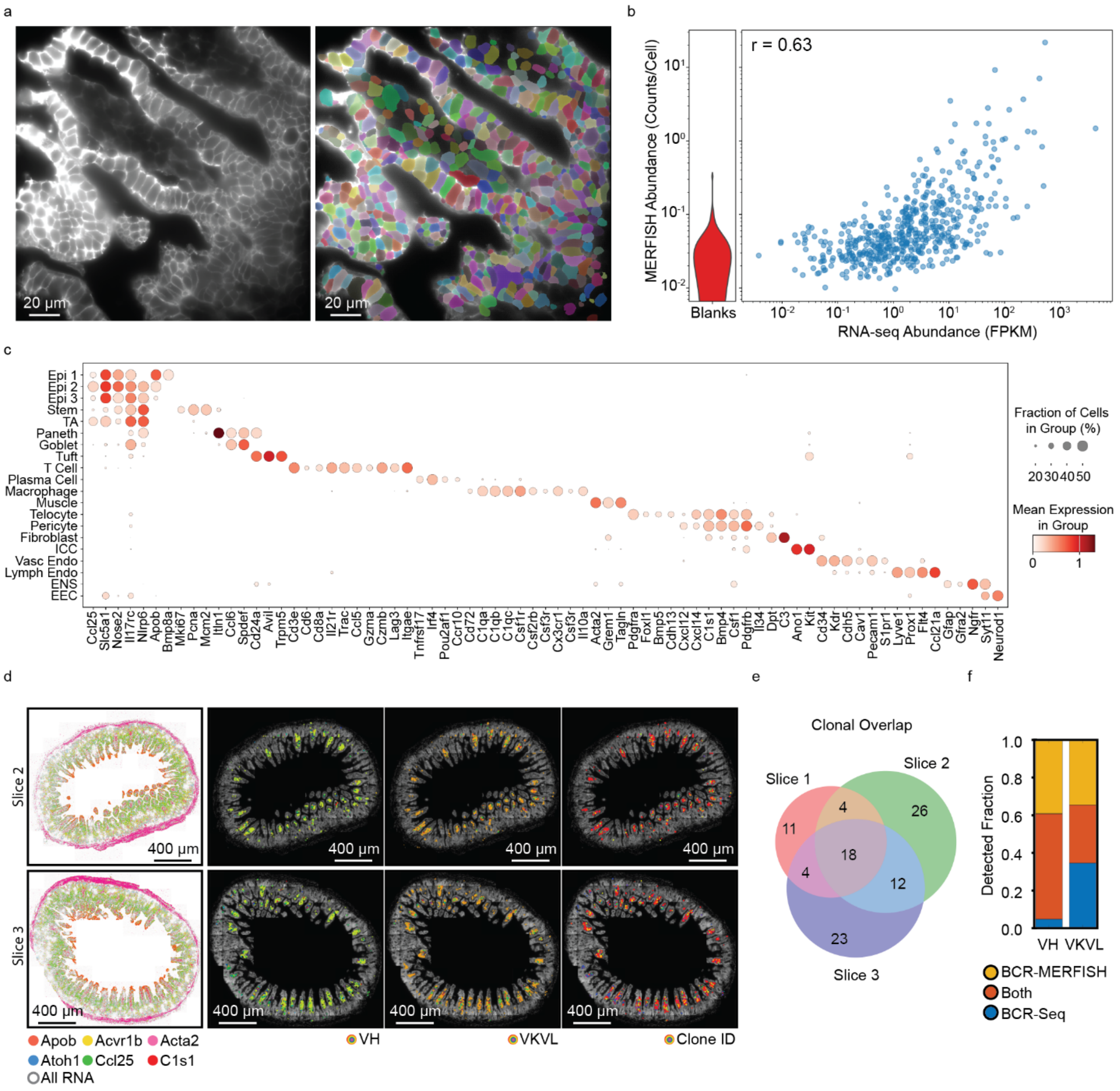


**Fig. S3 | Performance of BCR-MERFISH in the mouse ileum.** **a**, Representative image of a region of the mouse ileum stained with an antibody against a pan-cell-surface protein ($Na^{+}/K^{+}$-ATPase) with (right) and without (left) the cell masks (color) identified with Cellpose based on this stain. Scale bars: 20 µm. **b**, RNA abundance determined via MERFISH versus that determined by bulk RNA-sequencing for the 589 genes characterized with MERFISH (right) with the distribution of false-positive control barcodes (Blanks) determined via MERFISH (left) for the mouse ileum. r: Pearson correlation coefficient for the logarithmic expression. **c**, The average expression of key genes within clusters identified via MERFISH in the mouse ileum. Color indicates the average logarithmic expression while marker size indicates the fraction of cells with at least one copy of the RNA. **d**, The spatial distribution of all mRNAs (gray) and six example mRNAs (color) observed with MERFISH in two of the three slices used to compare BCR-MERFISH to BCR-seq (left) with the spatial distribution of all identified plasma cells plotted on top of the DAPI images collected for these slices (right) colored by the VH, VKVL, and clonal ID identified by BCR-MERFISH. Scale bars: 400 µm. **e**, Venn diagram of the overlap in identified plasma cell clones between the three ileal slices characterized with BCR-MERFISH. **f**, The fraction of all detected VH or VKVL genes with either BCR-MERFISH or BCR-seq that were detected in either technique individually or in both techniques.

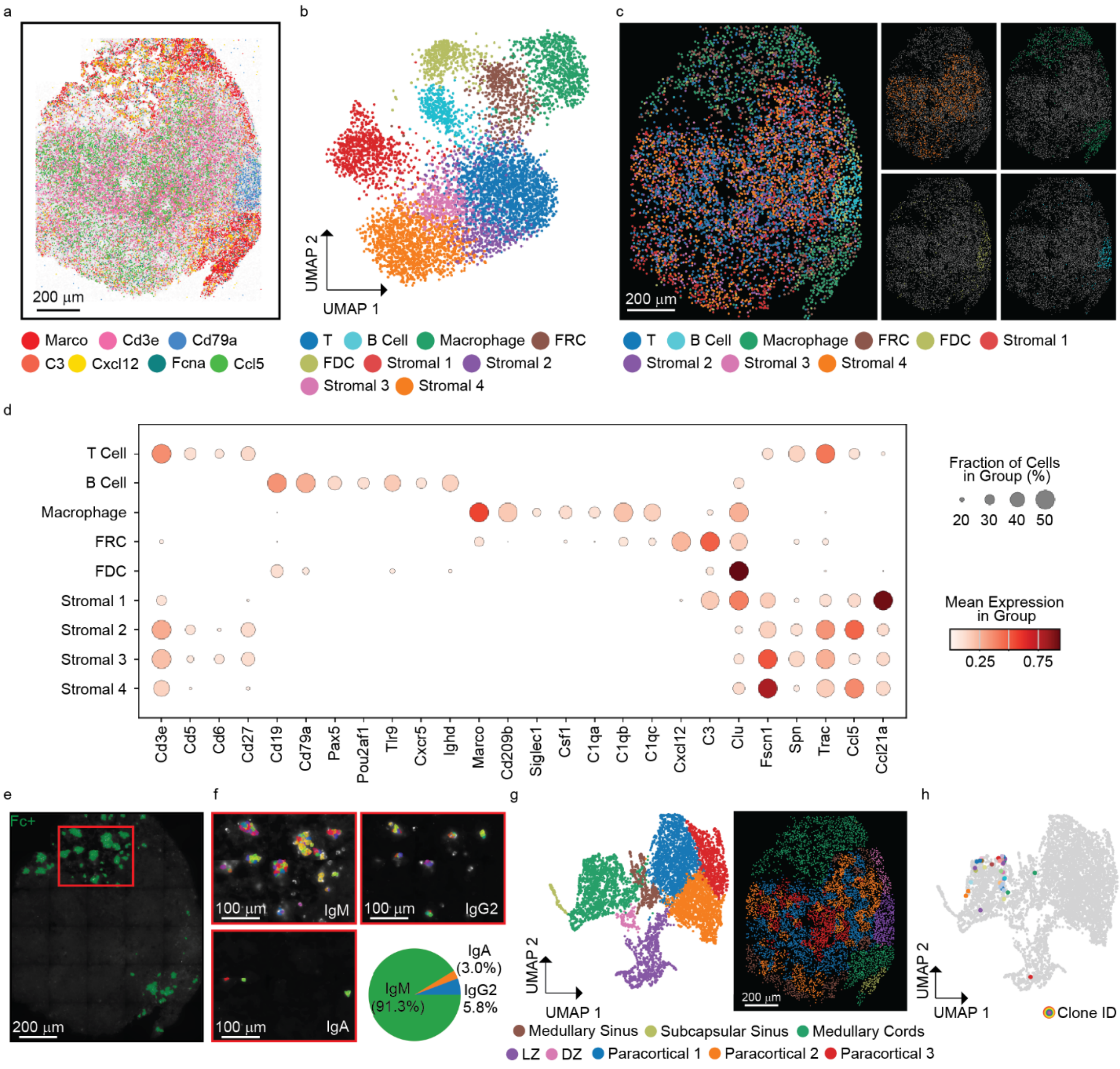


**Fig. S4 | BCR-MERFISH reveals plasma cell clonality and spatial and cellular context in a skin draining lymph node. a**, The spatial distribution of all mRNAs (gray) and seven example mRNAs (color) observed with MERFISH in a section of a mouse skin-draining lymph node. Scale bar: 200 µm. **b**, UMAP representation of the cells identified in the slice in (a) colored by cell type. **c**, The spatial distribution of the cells identified in the slice in (a) colored as in (b) for all cells (left) or individual cell types (right). Scale bar: 200 µm. **d**, The average expression of key genes within clusters identified via MERFISH in (b). Color indicates the average logarithmic expression while marker size indicates the fraction of cells with at least one copy of the RNA. **e**,**f**, Composite image of the slice in (a) combining all Fc stains (e) or for each Fc type with cells colored by clonal ID as determined via MERFISH (f). The pie chart represents the fraction of plasma cells associated with each Fc type. Scale bars: 200 or 100 µm. **g**, UMAP representation of the local cellular neighborhood colored by major anatomical features (left) and the spatial distribution of cells in this slice colored by neighborhood (right). Scale bar: 200 µm. **h**, Neighborhood UMAP as in (g) with all cells (gray) and plasma cells colored by the clonal ID determined via BCR-MERFISH.

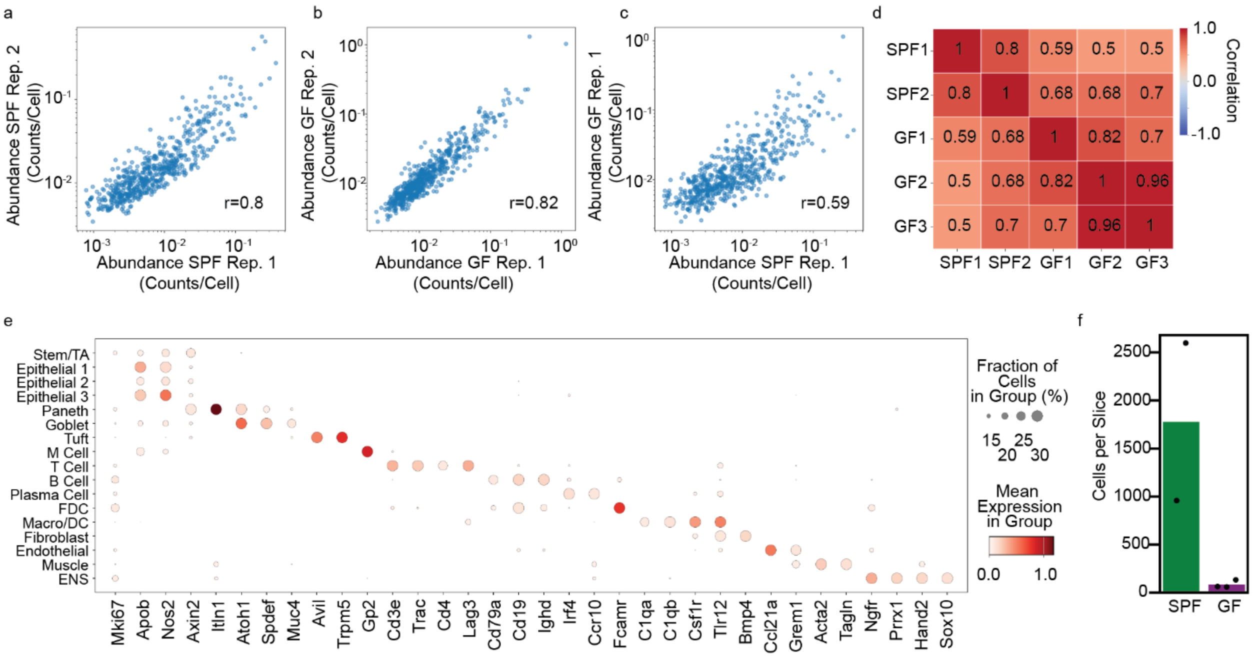


**Fig. S5 | Performance of MERFISH in the specific pathogen-free and germ-free ileum. a-c**, RNA abundance determined via MERFISH for one slice versus that determined for a different slice for two SPF mice (a), two GF mice (b), or an SPF and a GF mouse (c). r: Pearson correlation coefficient between the logarithmic expression. **d**, The pairwise Pearson correlation coefficients between RNA abundance determined via MERFISH for all SFP and GF mice. **e**, The average expression of key genes within clusters identified via MERFISH for all SPF and GF mouse ileum measurements. Color indicates the average logarithmic expression while marker size indicates the fraction of cells with at least one copy of the RNA. **f**, The number of plasma cells for each ileum slice from SPF or GF mice (markers). Bars represent the average over all measured mice of each condition.

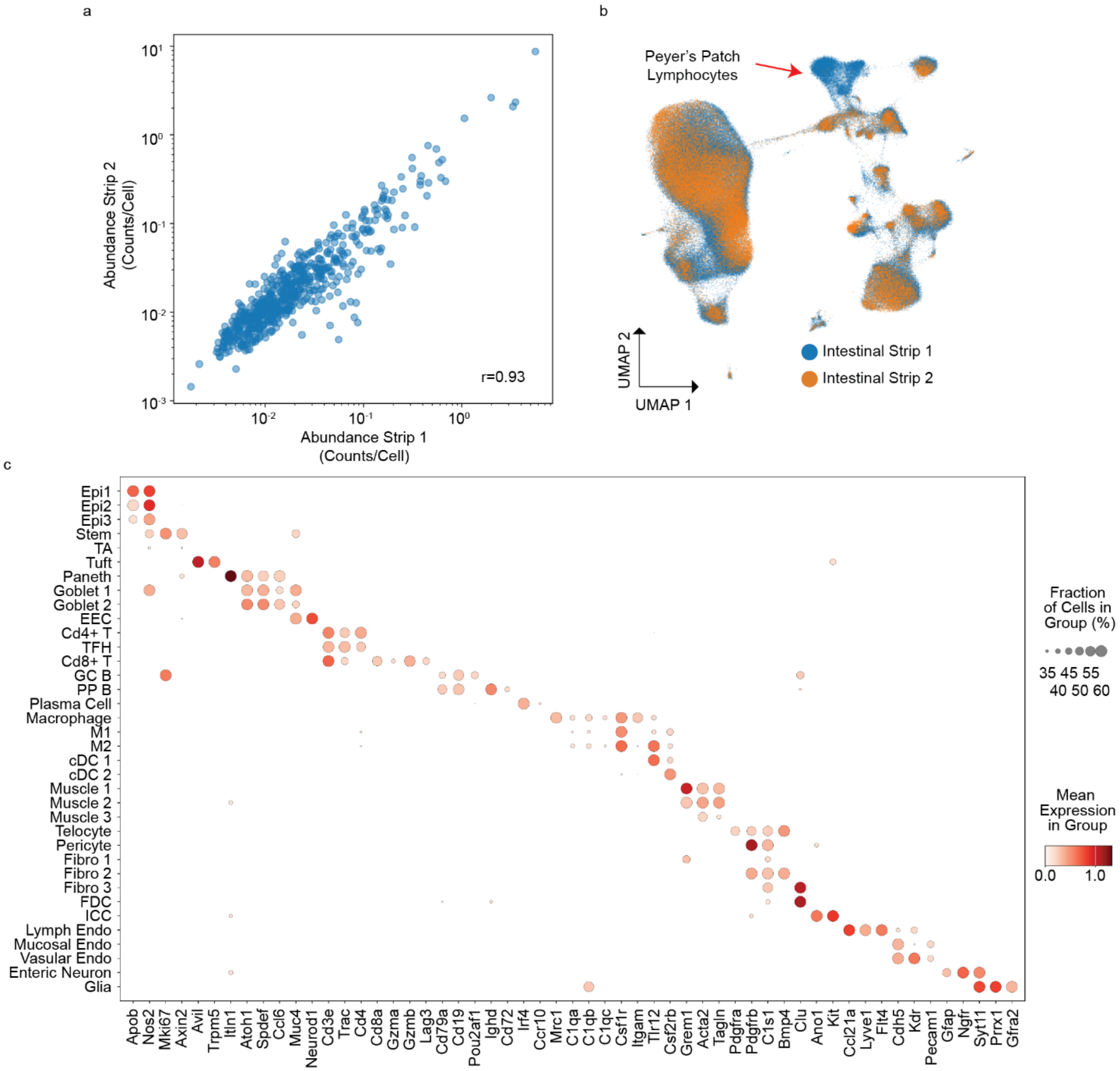


**Fig. S6 | Performance of MERFISH in ileal strips. a**, RNA abundance determined via MERFISH for each of the two intestinal strips. r: Pearson correlation coefficient between the logarithmic expression. **b**, UMAP representation of the cells observed in each of the two strips colored by strip or origin. Note that no batch correction was applied. Cells seen in only one replicate represent the PP seen in that strip but not the other and are highlighted. **c**, The average expression of key genes within clusters identified via MERFISH. Color indicates the average logarithmic expression while marker size indicates the fraction of cells with at least one copy of the RNA.

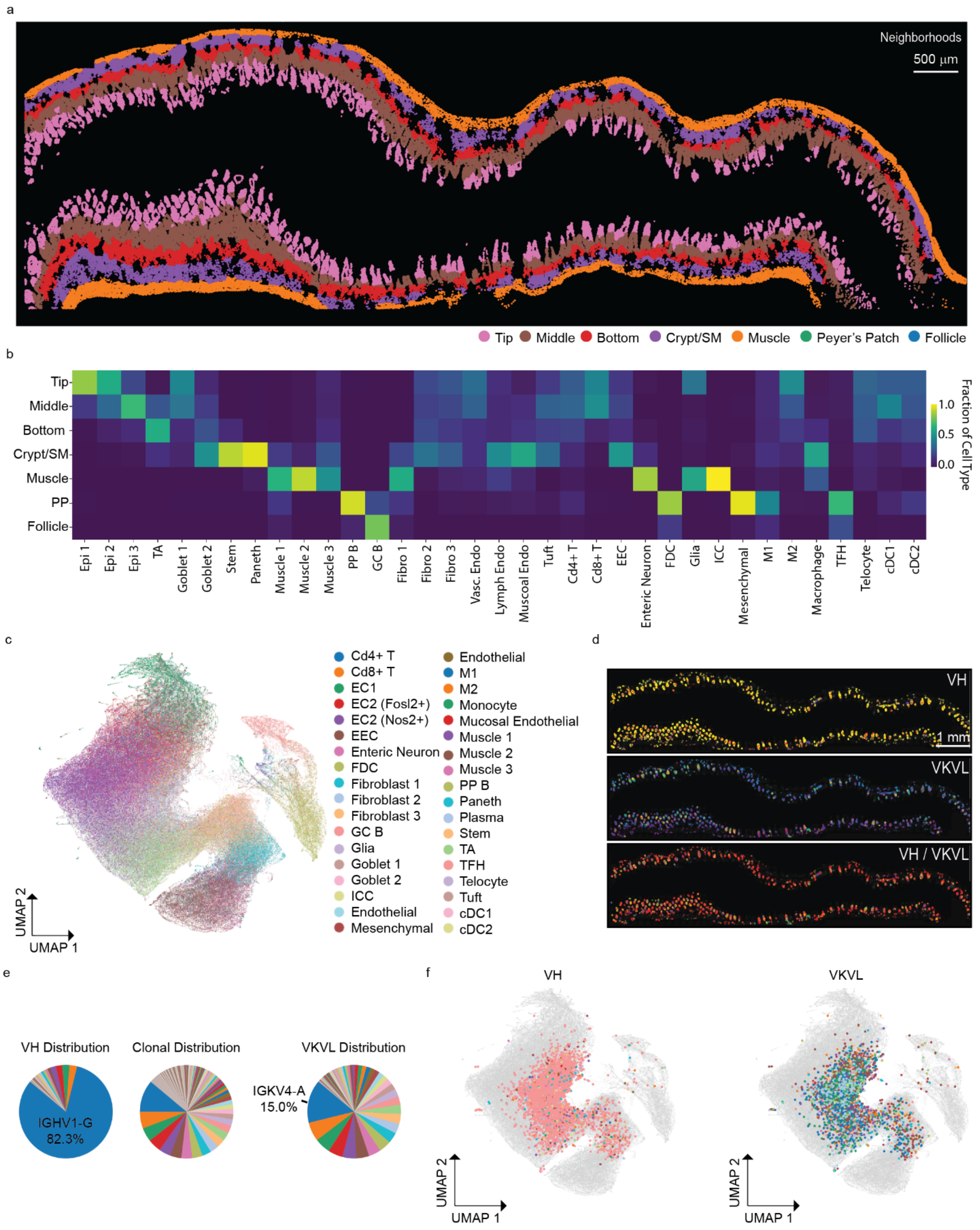


**Fig. S7 | The organization and distribution of plasma cell clones in ileal strips. a**, The spatial distribution of all cells in the second ileal strip colored by neighborhood assignment as in Fig. 4c.

Scale bar: 1 mm. **b**, The normalized fractional abundance of each cell type identified in the two intestinal strips within each identified neighborhood. **c**, UMAP representation of the neighborhoods as in Fig. 4c colored by cell types. **d**, The spatial distribution of plasma cells in the second ileal strip colored by VH (top), VKVL (middle), and clone ID (bottom) as determined by BCR-MERFISH. Scale bar: 1 mm. **e**, The fraction of plasma cells assigned a given VH gene (top left), a VKVL gene (bottom left), and a given clone ID (top right) for all measured plasma cells. **f**, UMAP representation of neighborhoods as in Fig. 4c with all cells (gray) and plasma cells colored the assigned VH (left) or VKVL (right) as determined by BCR-MERFISH.

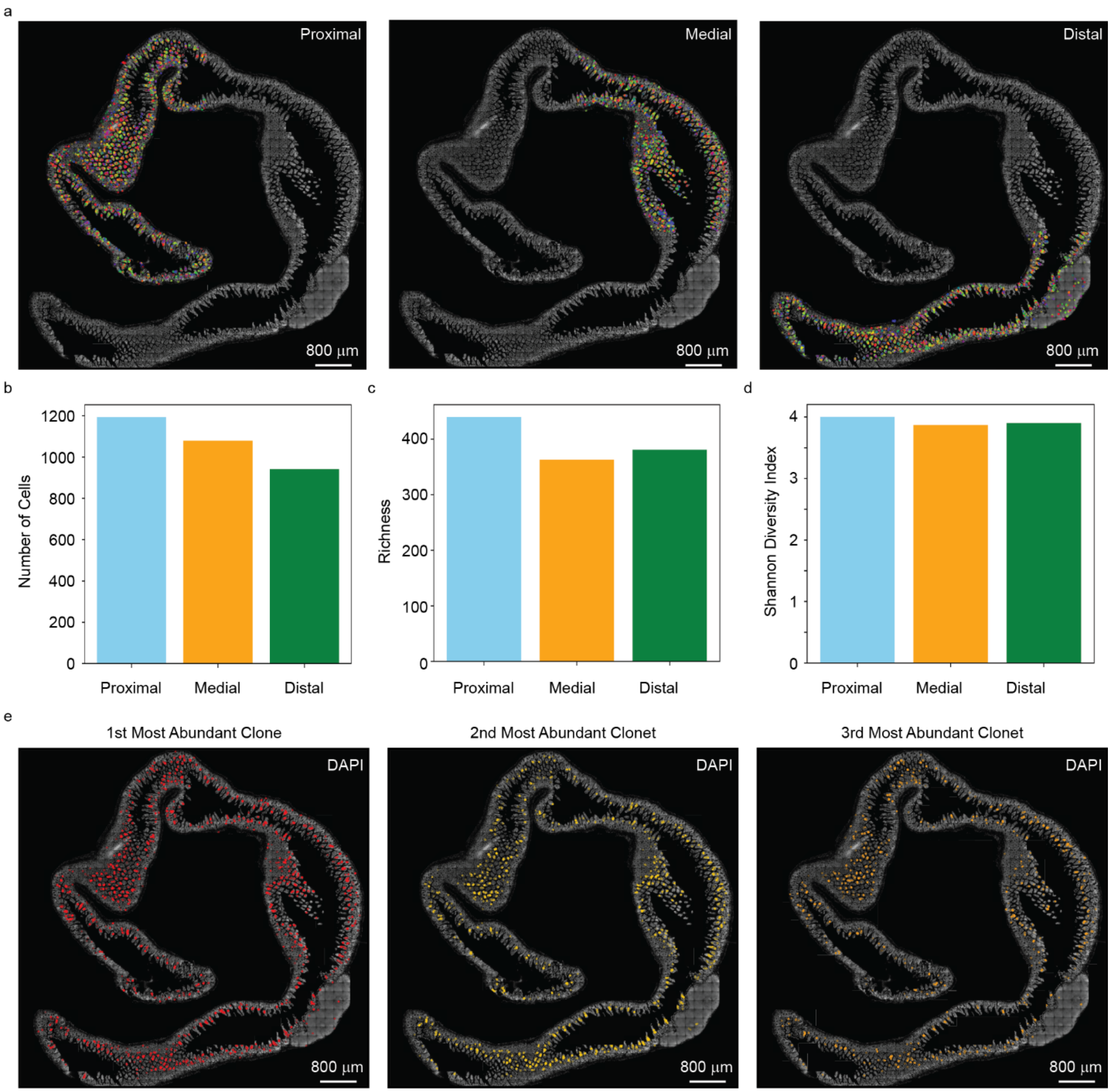


**Fig. S8 | Distribution of plasma cell clones in the first ileal strip. a**, DAPI image of the first ileal strip with the location of all plasma cells assigned to the proximal, medial, or distal ileal regions colored by the clone ID as determined by BCR-MERFISH. Scale bars: 800 µm. **b-d**, The total number of plasma cells (b), the richness in clonal IDs (c), and the Shannon Diversity Index for clonal IDs (d) for the proximal, medial, and distal ileal regions as in (a). **e**, DAPI image of the first ileal strip with the location of the top three most abundant plasma cell clones. Scale bars: 800 µm.

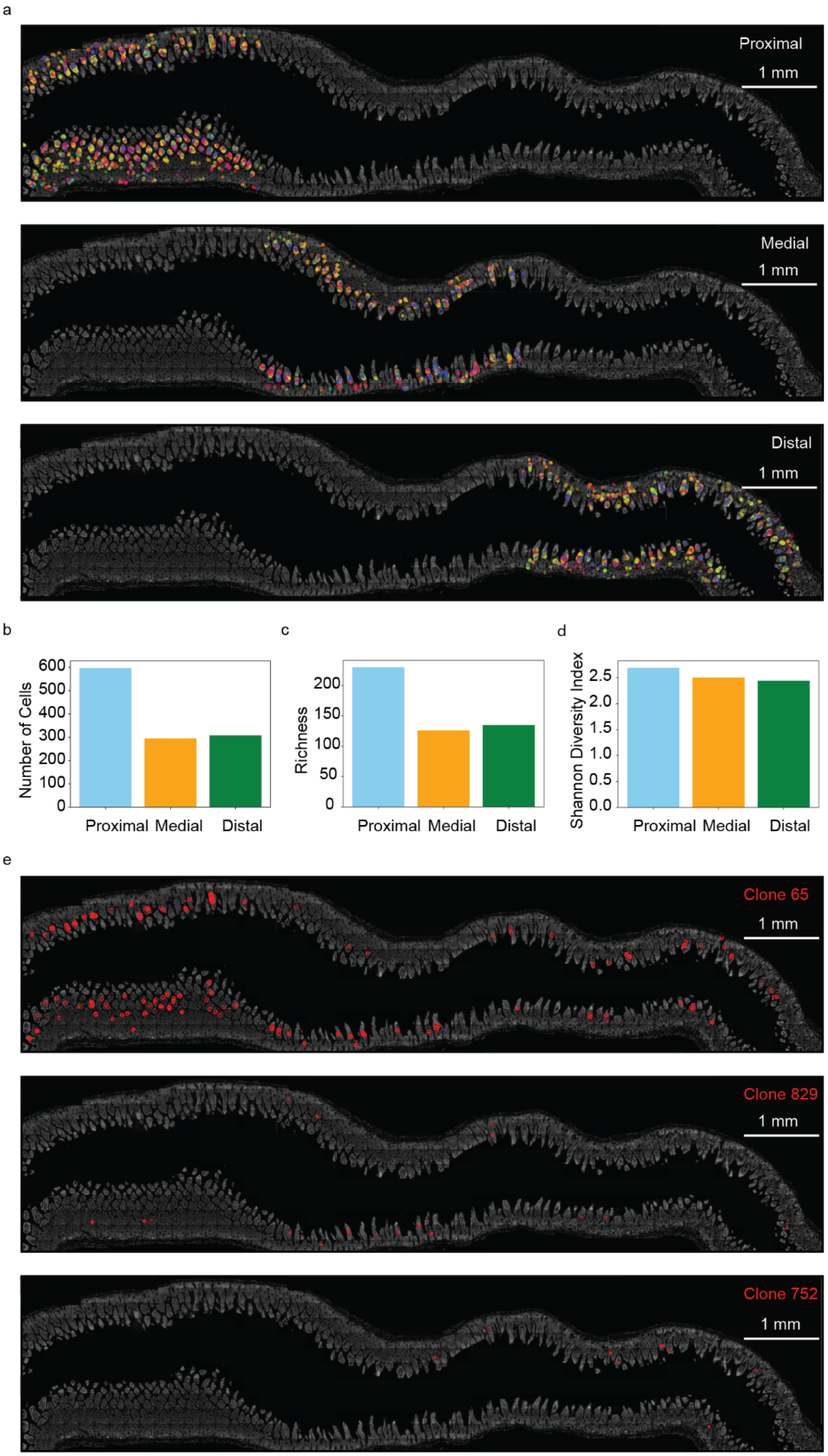
a
Proximal
1 mm
Medial
1 mm
Distal
1 mm
b
Number of Cells
600
500
400
300
200
100
0
Proximal
Medial
Distal
c
Richness
200
150
100
50
0
Proximal
Medial
Distal
d
Shannon Diversity Index
2.5
2.0
1.5
1.0
0.5
0.0
Proximal
Medial
Distal
e
Clone 65
1 mm
Clone 829
1 mm
Clone 752
1 mm

**Fig. S9 | Distribution of plasma cell clones in the second ileal strip. a**, DAPI image of the second ileal strip with the location of all plasma cells assigned to the proximal, medial, or distal ileal regions colored by the clone ID as determined by BCR-MERFISH. Scale bars: 1 mm. **b-d**, The total number of plasma cells (b), the richness in clonal IDs (c), and the Shannon Diversity Index for clonal IDs (d) for the proximal, medial, and distal ileal regions as in (a). **e**, DAPI image of the second ileal strip with the location of the three plasma cell clones that showed proximal (top), medial (middle), or distal (bottom) enrichment. Scale bars: 1 mm.

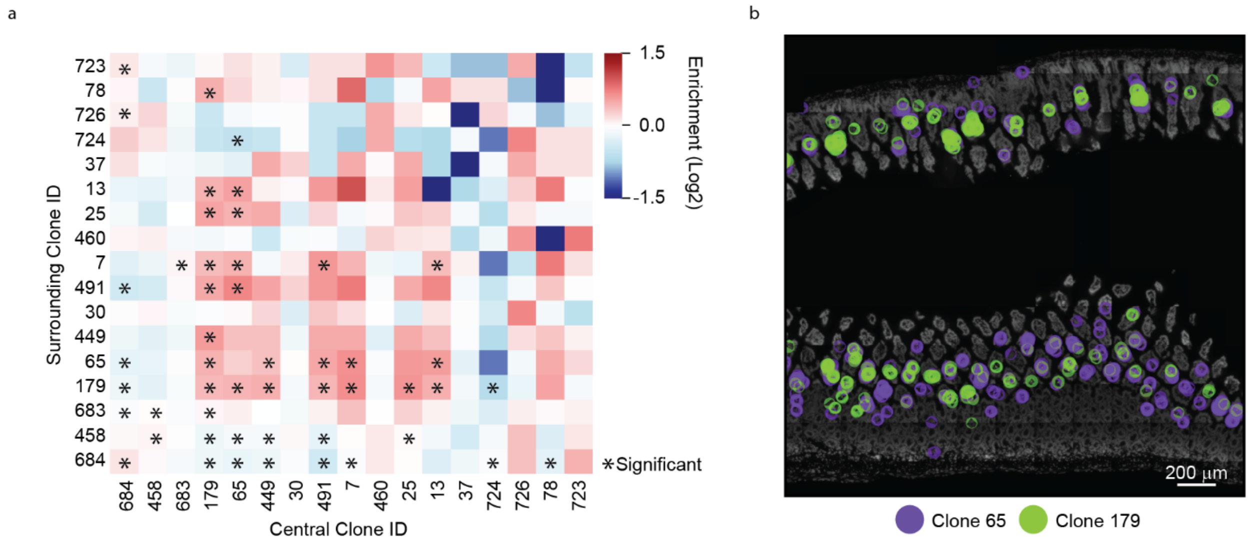


**Fig. S10 | Spatial Distribution of Plasma Cell Clones that Spatially Co-Occur in the Second Intestinal Strip. a**, The enrichment in the spatial co-occurrence for plasma cells of one clonal type versus those of another for the second ileal strip. Clone IDs are sorted by abundance in descending order. Columns represent the central clone while rows represent the clones in the surrounding neighborhood. **b**, DAPI image of the representative locations within a second ileal strip with the location of the listed plasma cell clones plotted. Scale bars: 200 µm.

### Supplementary Tables and Captions

**Supplementary Table 1. Grouping of VH and VKVL genes and barcode assignment (provided as a separate xlsx file).** The first sheet ('VH') contains information regarding VH gene groupings while the second sheet ('VKVL') contains information regarding VKVL gene groupings. In each sheet, the column 'Node Name' is the name given to the V gene group created from our homology-aware probe design pipeline, the column 'Genes Included in Node' lists the IGHV, IGKV, or IGLV genes included in each node, and the column 'Barcode' lists the binary barcode assigned to each node. The sheets entitled 'VH Collision' or 'VKVL Collision' contain information regarding the collision barcodes associated VH or VKVL genes. The column 'Name' lists the name of the collision, and the column 'Barcode' lists the barcode associated with that collision.

**Supplementary Table 2. Oligonucleotide sequences for BCR-MERFISH (provided as a separate xlsx file).** The sheets titled 'VH', 'VKVL', or 'Fc' contain the name and sequence associated with all encoding probes (Fc) or encoding probe template molecules (VH and VKVL) encoding the VH gene groups, the VKVL gene groups, and the constant regions. The sheet entitled '564Igi' contain the smFISH probe sequences targeting the V genes associated with the 564Igi mouse. The properties of all readout sequences are in the sheet entitled 'Readout Sequences'. In this sheet, the 'Name' column contains a name for each readout probe, the 'Sequence' column contains its sequence, the 'Library' column contains the specific target of the encoding probes for which that readout was used, and the 'Bit' column contains the location in the barcode associated for each library. 'Cy5-S-S-' indicates conjugation of a readout sequence to a Cy5 fluorophore via a disulfide bond while 'Alexa750-S-S-' indicates similar conjugation to an Alexa750 fluorophore.

**Supplementary Table 3. Codebook for 589-gene MERFISH panel (provided as a separate xlsx file).** The column 'Name' contains the name of the targeted gene, the column 'ID' contains a unique isoform ID associated with the targeted gene, and the column 'Barcode' contains the binary barcode associated with each gene. The readout probe sequences associated with each bit in these barcodes are contained in Table S2. Entries with a name containing 'Blank' are barcodes not assigned to an RNA and which, thus, serve as false positive controls.

**Supplementary Table 4. Encoding probe template sequences for 589-gene MERFISH panel (provided as a separate xlsx file).** The column 'Name' contains a name for the template molecule and the column 'Sequence' contains the sequence of the template molecule used to make the corresponding MERFISH encoding probe.

**Supplementary Table 5: Clonal identity and abundance in all BCR-MERFISH measurements (provided as a separate xlsx file).** Each sheet lists the clonal identity and abundance for plasma cells measured across each replicate for all BCR-MERFISH experiments. The sheet entitled 'BCR-seq' lists the measures used to compare to 'BCR-Seq'. The sheet entitled 'SPF vs GF' lists the results for the comparison between SFP and GF mice. The sheet entitled 'Lymph Node' lists the results in the skin draining lymph node. The sheet entitled 'Intestinal Strips' lists the results in the ileal strips. Within each sheet the column 'Clone_ID' lists the name of the clone; the columns 'Unique_Pair' lists the VH and VKVL genes associated with the clone; the 'Total Count' column lists the number of cells associated with that Clone ID across all replicates; and all additional columns list the number of each of a given clone observed in the replicate named in that column.

**Supplementary Table 6. Previously reported public VH and VK genes (provided as a separate xlsx file).** The 'Public Gene' column represents the gene that was reported as a public V gene by previous works[26,45]. The 'BCR-MERFISH Group' column represents the BCR-MERFISH probe group that was contains that gene. The 'Putative Public Clones' column lists V genes observed across multiple mice with BCR-MERFISH but which were not reported previously.

**Supplementary Table 7. Statistical tests for neighborhood enrichment for plasma cell clones (provided as a separate xlsx file).** The 'Clone_ID' column contains a unique name for each clone. The 'Unique_Pair' column represents the VH and VKVL pair that constitutes the Clone ID. The 'Chi2' column lists the chi-squared value that the clone is enriched in any given neighborhood. The 'P-value' column lists the p-value associated with the chi-squared test while the 'P-value_corrected_BH' and 'Significant_BH' columns list the Benjamini-Hochberg corrected p-value and whether that value passes a significant threshold, after correction, of 0.05. The columns named with neighborhood names as in Fig. 4 represent the fraction of each clone in each neighborhood. Entries are sorted by p-value. Both replicates are listed in the same sheet.

**Supplementary Table 8. Statistical tests for proximal-distal enrichment for plasma cell clones (provided as a separate xlsx file).** The 'Clone_ID' column contains a unique name for each clone. The 'Unique_Pair' column represents the VH and VKVL pair that constitutes the Clone ID. The 'Chi2' column lists the chi-squared value that the clone is enriched in any ileal region. The 'P-value' column lists the p-value associated with the chi-squared test while the 'P-value_corrected_BH' and 'Significant_BH' columns list the Benjamini-Hochberg corrected p-value and whether that value passes a significant threshold, after correction, of 0.05. The columns named with region names as in Fig. 4 represent the fraction of each clone in each region. The single sheet includes both replicates.

**Supplementary Table 9. Statistical tests for local spatial enrichment of plasma cell clones (provided as a separate xlsx file).** The columns ‘Clone_ID1’, and ‘Clone_ID2’ represent the pair of clones considered. The ‘Z-score’ column contains the observed enrichment scaled by the mean and the standard deviation of that enrichment determined by resampling. The ‘P-value’ column contains the significance of this z-score using a two-sided t-test. The ‘P-value_corrected_BH’ contains adjusted p-values after Benjamini-Hochberg corrections. The ‘Significant_BH’ column is true if this adjusted p-value is less than 0.05. Both replicates were treated separately, and the values are provided in two different sheets.
